# Scaling laws of plasmids across the microbial tree of life

**DOI:** 10.1101/2024.10.04.616653

**Authors:** Rohan Maddamsetti, Maggie L. Wilson, Hye-In Son, Zhengqing Zhou, Jia Lu, Lingchong You

## Abstract

Plasmids play a critical role in shaping the dynamics and evolution of microbial communities. The capacity of a plasmid to express genes is constrained by two parameters: length and copy number. However, the interplay between these parameters and their constraints on plasmid evolution have remained elusive due to the absence of comprehensive quantitative analyses. To address this gap, we developed Probabilistic Iterative Read Assignment (PIRA), a new computational method that overcomes previous computational bottlenecks, enabling rapid and accurate determination of plasmid copy numbers at an unprecedented scale. Applying PIRA to all microbial genomes in the NCBI RefSeq database with linked short-read sequencing data in the Sequencing Read Archive (SRA), we analyzed 4,317 bacterial and archaeal genomes encompassing 11,338 plasmids, spanning the microbial tree of life. Our analysis reveals three scaling laws of plasmids: first, an inverse power-law correlation between plasmid copy number and plasmid length; second, a positive linear correlation between protein-coding genes and plasmid length; and third, a positive correlation between metabolic genes per plasmid and plasmid length, particularly for large plasmids. These scaling laws imply fundamental constraints on plasmid evolution and functional organization, indicating that as plasmids increase in length, they converge toward chromosomal characteristics in copy number and functional content. Our findings not only advance the understanding of plasmid dynamics but also have implications for microbial evolution, biotechnology, and the design of synthetic plasmids.

**Significance:** By discovering universal scaling laws and developing a new computational method to compute plasmid copy numbers across the microbial tree of life, we show that as plasmids increase in length, they converge to chromosomes in their copy number and their coding and metabolic properties. This insight reveals fundamental principles governing plasmid evolution and has implications for biotechnology and medicine.

## Introduction

Plasmids are extrachromosomal DNA elements, found ubiquitously across bacteria and archaea, that mediate the flow of genes within and across microbial communities. Plasmids play a role in how microbial populations rapidly adapt to novel selection pressures, by amplifying the copy number of beneficial genes and promoting their spread by horizontal gene transfer^1–3^. In the context of human health, plasmids shape human microbiome dynamics^4–6^, in particular by serving as critical vectors for the dissemination of antibiotic resistance^7^. Plasmids are also foundational to biotechnology, as they can engineered to control recombinant gene expression and the behavior of cells, populations, and microbial consortia^8,9^.

The capacity of a plasmid to express genes is determined by its length and by its copy number. Here we define the plasmid copy number (PCN) as the number of plasmid copies per the longest chromosome per cell. Understanding the interplay between plasmid length and PCN is essential, as these parameters^10^ affect the molecular biology, ecology, and evolution of microbes. Such understanding also has applications for engineering microbial populations and communities^11^.

Intuitively, we would expect that plasmid lengths and copy numbers constrain each other. For instance, a cell may have a limited capacity to accommodate additional genetic material beyond the chromosome, which would impose a tradeoff between PCN and length. If so, then increasing PCN would necessarily constrain the functional capacity of a plasmid, in terms of the number and functional diversity of the genes it carries. Small-scale data supports this hypothesis: high-copy-number plasmids are often small^6^, while larger conjugative plasmids often have 1-2 copies per cell^1^.

However, comprehensive data on the distribution of PCN across the microbial tree of life do not exist, and whether and how it is correlated to plasmid length remains elusive. The rapidly increasing amount of sequence data on plasmid-bearing microbes creates an opportunity to address these questions. A major technical challenge is that direct PCN calculations at scale require pairwise sequence alignment between thousands of sequencing datasets and reference genomes, a process that is computationally prohibitive^12–18^. The computational costs associated with sequence alignment thus represent a key bottleneck that restricts plasmid copy number computations from approaching the scope or scale of all microbial genomes.

To overcome this bottleneck, we developed Probabilistic Iterative Read Assignment (PIRA), which uses pseudoalignment to rapidly and accurately estimate plasmid copy numbers across large datasets. By applying PIRA to all complete genomes containing plasmids in the NCBI RefSeq database^19^ with linked short-read sequencing data in the Sequencing Read Archive (SRA)^20^, we report the largest dataset on plasmid lengths and copy numbers to date. Our analysis encompasses 4,317 bacterial and archaeal genomes and 11,338 plasmids, spanning the microbial tree of life. We discovered universal scaling laws governing plasmid biology: first, an inverse power-law correlation between PCN and plasmid length; second, a positive linear correlation between protein-coding genes and plasmid length; and third, a positive correlation between metabolic genes per plasmid and plasmid length, particularly for large plasmids. These scaling laws imply fundamental constraints on the evolution of plasmids, as well as their ability to accommodate functional traits. Our findings reveal that as plasmids increase in length, they converge toward chromosomal characteristics in copy number and functional content, challenging traditional distinctions between plasmids and chromosomes. This discovery not only advances our understanding of plasmid dynamics and microbial evolution but also has implications for biotechnology, such as the rational design of synthetic plasmids and the engineering of microbial communities.

## Results

### PCNs are rarely reported in the microbial genomics literature

One goal of this project was to generate a comprehensive dataset of PCNs, because we found that few plasmids had reported copy numbers in the literature. To quantitatively assess this research gap, we randomly sampled 50 genomes, each containing at least one multicopy plasmid with PCN > 10 (Methods). We manually examined the publications associated with each genome, based on the genome annotation files found in the NCBI RefSeq database. PCNs were reported for 3 out of 50 genomes (Supplementary Table 1). Therefore, we estimate that ∼6% of genomes with sequenced plasmids have PCNs that are reported in the literature.

### A scalable bioinformatics pipeline for PCN estimation

To address this critical gap and enable a comprehensive understanding of plasmid dynamics across the microbial tree of life, we built a scalable bioinformatics pipeline (Figure 1A and Supplementary Data 1). Using it, we estimated PCNs at an unprecedented scale, facilitating the discovery of universal scaling laws in plasmid biology. Specifically, 19,538 bacterial and archaeal genomes containing plasmids in the NCBI RefSeq database were screened for linked short read sequencing data in the Sequencing Read Archive (SRA). Short-read sequencing data for 4,540 genomes containing plasmids were downloaded, and the sequencing reads were successfully pseudoaligned to chromosomal and plasmid reference sequences in 4,510 genomes. Plasmids with fewer than 10,000 mapped reads were removed from the analysis. Our final dataset comprises 4,317 bacterial and archaeal genomes with a total of 11,338 PCN estimates. Pseudoalignment overcomes the computational bottleneck that would be caused by using traditional alignment to map terabytes worth of short-read sequencing data to thousands of reference microbial genomes^12–18,21,22^.

**Figure 1.**
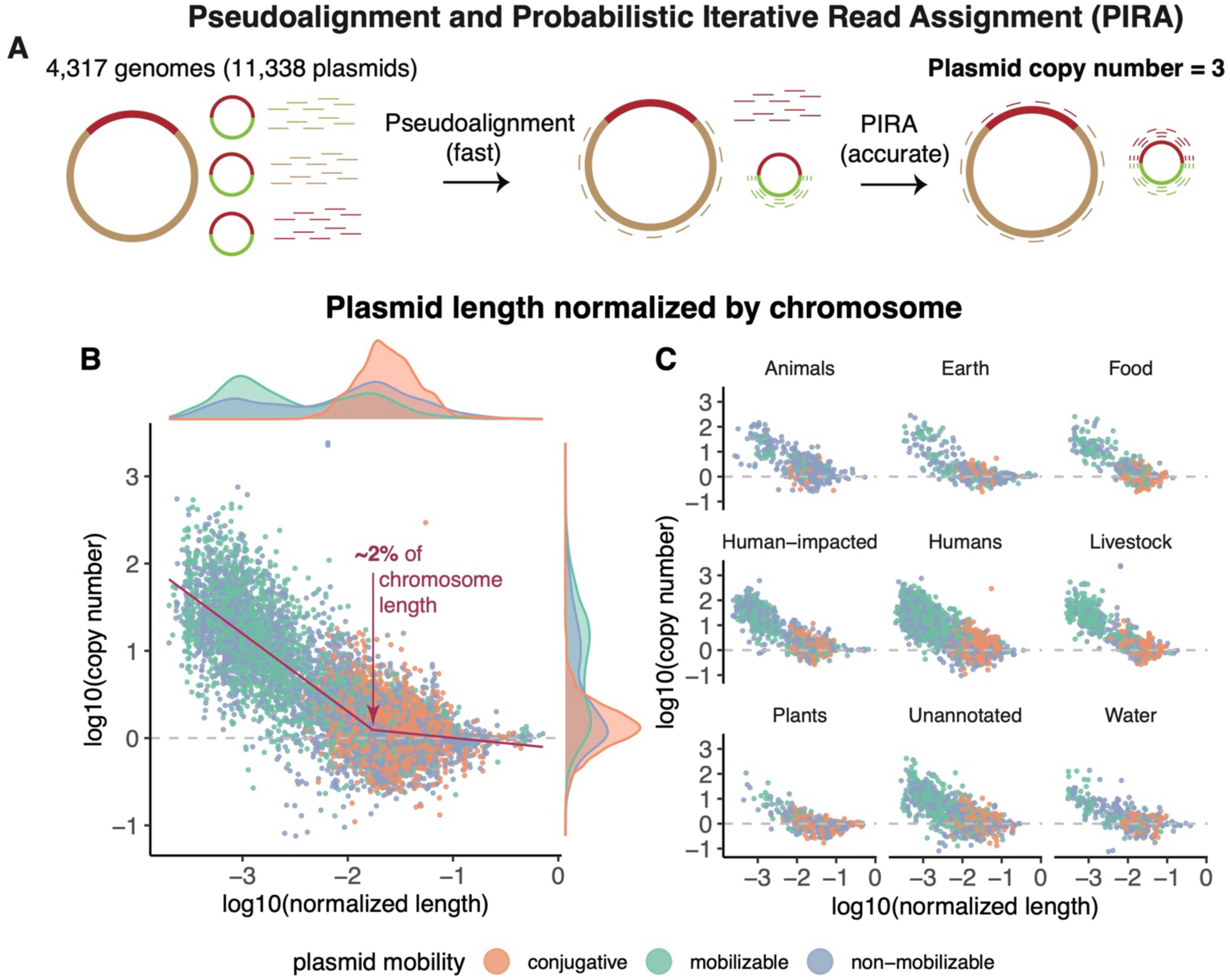
Computation of plasmid copy numbers over the microbial tree of life reveals that copy number inversely correlates with plasmid length. **A)** The computational pipeline. Suppose a genome has one chromosome and one plasmid with 3 copies relative to the chromosome. By dividing the total amount of sequencing data mapped to chromosome or plasmid by the length of the chromosome and plasmid, the ratio of plasmid DNA to chromosomal DNA can be calculated. This estimates the plasmid copy number per chromosome in the sample. Pseudoalignment is used to rapidly estimate plasmid copy numbers, and Probabilistic Iterative Read Assignment (PIRA) is used to incorporate reads that map to multiple replicons (e.g., the chromosome and plasmid) to further improve plasmid copy number estimates. **B)** Plasmid length inversely correlates with plasmid copy number. Rescaling plasmid length by the length of the largest chromosome in the cell reveals a scaling law. A segmented regression (in maroon) was fit to these normalized data on a log-log plot. The segmented regression has a first slope of –0.89, a breakpoint at –1.76 (or 1.74% of a chromosome), a second slope of –0.12, and an Adjusted R2 of 0.691. The marginal density distributions of plasmid copy number and normalized length are displayed on the axes. **C)** The inverse correlation holds across diverse environments. The ecological provenance of each replicon was annotated per the method described in Maddamsetti et al.^3^ (Methods).

Our PCN calculation only depends on two assumptions. First, we assume that the lengths of all chromosomes and plasmids in the genome are known. Second, we assume that the relative amounts of sequencing data that map to chromosomes versus plasmids in each sequencing sample is proportional to the physical amount of DNA corresponding to chromosomes and plasmids in the genome.

Figure 1A shows an example calculation. Suppose a genome has one chromosome and one plasmid with 3 copies relative to the chromosome. By dividing the total amount of sequencing data (in units of nucleotide base pairs) mapped to chromosome or plasmid by the corresponding lengths of the chromosome or plasmid, the ratio of plasmid DNA to chromosomal DNA in the sequencing data can be calculated. This estimates the PCN per chromosome in the sequenced sample.

### PIRA accounts for sequencing reads that map to multiple replicons

The direct PCN estimation method above does not take sequencing reads that map to multiple replicons into account. Here, we define “replicon” as a generic term for either chromosomes or plasmids. We define a “uniread” as a sequencing read that unambiguously maps to a single replicon, and a “multiread” as a sequencing read that maps to multiple replicons. Multireads can arise due to repetitive or duplicated sequences that are shared across replicons. Such a situation can arise when plasmids and chromosomes share mobile genetic elements, such as a transposon that has jumped from the chromosome to a plasmid^2,3^. Multireads may affect PCN estimates when a plasmid shares significant homology with either the chromosome, or other plasmids in the cell. In this case, an unknown fraction of multireads may come from the plasmid of interest, while the remainder comes from other replicons in the genome. However, that unknown fraction depends on the PCN, introducing a circular dependency.

To solve the multiread problem and more accurately estimate PCNs, we developed Probabilistic Iterative Read Assignment (PIRA). Pseudoalignment is first used to map reads to replicons.

Unireads are used to make an initial estimate of PCNs. The multireads are then re-aligned to the reference genome using traditional pairwise sequence alignment^23,24^. The multireads that map to a single genomic location with traditional alignment are combined with the unireads to improve the PCN estimates (Supplementary Data 1). The remaining multireads are then probabilistically allocated to each replicon in the genome, based on the initial PCN estimates. The estimates are iteratively updated until convergence, based on the re-allocation of multireads. That is, our pipeline therefore uses pseudoalignment to quickly make good initial PCN estimates and then uses PIRA to refine those estimates based on multiread information.

### PIRA accurately estimates PCN

We used PIRA to estimate the copy numbers of 11,381 plasmids (Supplementary Data 2). By comparing PCN estimates made with PIRA to PCN estimates by the direct method, we find that PIRA enables 1,563 more estimates, by enabling estimation for plasmids with many multireads but few unireads. PIRA also improves the accuracy of PCN estimates made by the direct approach, which neglects multireads (Supplementary Figure S1A).

Several plasmids had estimated PCN < 1, meaning that the number of plasmid copies were lower than the number of chromosome copies in the sample sent for genome sequencing. To assess PIRA’s accuracy, we first compared PCN estimates generated by PIRA to PCN estimates generated by traditional alignment algorithms^23–25^. Due to the computational overhead of using alignment to estimate PCNs, we selected a random set of 100 genomes to benchmark PIRA against traditional alignment methods. Each of these randomly selected 100 genomes contained at least one plasmid with an estimated PCN < 0.8, to test whether these low PCN estimates were also recovered by traditional alignment algorithms. If so, this outcome would indicate that these low PCN estimates were a property of the underlying sequencing data, and not an artifact caused by PIRA.

We used two methods to estimate PCN using traditional alignment. First, we used minimap2, as a state-of-the-art method for pairwise alignment of sequencing reads to reference chromosomes and plasmids^23,24^. Second, we used breseq^25^, an established genome resequencing pipeline which which uses Bowtie 2^26^ to align sequencing reads to reference chromosomes and plasmids. Importantly, both minimap2 and breseq have been used to estimate PCNs^2,3,27,28^.

The comparison between PIRA and minimap2 (Supplementary Figure S1B) and between PIRA and breseq (Supplementary Figure S1C) shows that PCN estimates generated by PIRA are consistent with PCN estimates generated by traditional read alignment algorithms. Together, these results indicate that PIRA provides accurate and reliable PCN estimates and show that the low PCN estimates are a property of the underlying sequencing data, given the consistency of these estimates across methods.

As an additional technical control, we examined whether PCN estimates generated by pseudoalignment were sensitive to the specific choice of software used. We compared PCN estimates generated by Themisto^22^ (the software used for PIRA) against PCN estimates generated by kallisto^13^. As expected, we found that these PCN estimates were highly consistent (Supplementary Figure S1D).

### A universal inverse power-law correlation between plasmid length and PCN

Using PIRA, we generated the largest dataset on PCNs to date, covering bacterial and archaeal genomes across the microbial tree of life. This dataset reveals an inverse power-law correlation between PCN and plasmid length (Figure 1B and Supplementary Figure S2). The distribution of plasmid sizes is bimodal, so K-means clustering with K = 2 was used to assign plasmids into two clusters. The cluster of small plasmids has a mean copy number of 28.4 plasmids per chromosome, and a mean length of 6,433 bp. The cluster of large plasmids has a mean copy number of 1.79 plasmids per chromosome, and a mean length of 137,704 bp. MOB-typer^29^ was used to classify plasmids as conjugative, mobilizable, or non-mobilizable. 3,103 out of 3,126 conjugative plasmids fall into the cluster of large plasmids (99.3%). By contrast, mobilizable plasmids (i.e., those that can be transferred by conjugation, but do not themselves encode conjugation machinery) and non-mobilizable plasmids may be either large or small (Figure 1B). Several plasmids are longer than 500,000 bp in length. Among these megaplasmids^30^, 29 are conjugative, 28 are mobilizable, and 101 are non-mobilizable, consistent with previous observations that most megaplasmids are non-mobilizable^31^. Many of these megaplasmids are chromids, which resemble secondary chromosomes even though they replicate using plasmid replication and partitioning systems^30,32^.

This inverse correlation between PCN and plasmid length is universal, holding across diverse environments (Figure 1C and Supplementary Figure S2) and microbial taxa across the tree of life (Supplementary Figure S3). Furthermore, this inverse correlation between PCN and length largely holds within individual genomes as well. Out of 2,629 genomes containing two or more plasmids, 2,160 have an inverse correlation between plasmid length and copy number (mean Pearson correlation coefficient = –0.91), while 469 show a positive correlation (mean Pearson correlation coefficient = 0.85). Out of 1,615 genomes containing three or more plasmids, 1,445 have an inverse correlation between plasmid length and copy number (mean Pearson correlation coefficient = –0.87), while 170 show a positive correlation (mean Pearson correlation coefficient = 0.60).

We normalized the length of each plasmid by the length of the longest chromosome in its genome (Figure 1B and 1C). These data are well fit by a segmented regression model^33,34^ in which PCN linearly scales with plasmid length on a log-log scale, up to a length threshold at which PCNs start to converge to chromosomal copy numbers (Figure 1B and Supplementary Figure S2). The breakpoint for this scaling law occurs when plasmid reaches 1.7% of the length of the chromosome. A model comparison using Akaike’s Information Criterion shows that the segmented regression model (AIC = 7,803.4) is significantly better than both a linear regression model (AIC = 9,228.8) and a second-order polynomial regression model (AIC = 8,003.2). The same pattern holds when a segmented regression model is fit to the unnormalized data: the segmented regression (AIC = 8,757.4) is significantly better than both a linear regression model (AIC = 10,141.7) and a second-order polynomial regression model (AIC = 8,972.3). The breakpoint for the segmented regression fit to the unnormalized length data occurs at 56,234 bp.

The segmented regression model suggests the following interpretation. PCNs can be modeled as a mixture of plasmids with cell-cycle dependent and cell-cycle independent replication mechanisms. The low-copy number conjugative F plasmid (length 99,159 bp) has 1-2 copies per cells, and replicates in sync with the cell cycle in *Escherichia coli*^35–37^. By contrast, the multicopy R6K plasmid (39,872 bp) replicates in a cell-cycle independent manner^36,38,39^. The scaling law may represent how PCN scales with plasmid length for plasmids that replicate using cell-cycle independent mechanisms. Once plasmids reach a critical length threshold, which seems to be ∼2% of the length of the chromosome, mechanisms that coordinate plasmid replication with cell division become critical for stable plasmid maintenance.

### Small multi-copy plasmids mostly coexist with large low-copy plasmids

Plasmids often co-occur with other plasmids in the environment^40^, and positive interactions between plasmids can stabilize such co-existence within cells^41^. We asked how often multi-copy plasmids co- existed with larger plasmids in these data. Out of 4,317 genomes containing plasmids with PCN estimates, 1,688 contain a single plasmid. By contrast, out of 1,314 genomes containing plasmids with PCN > 10 in our data, only 159 contained those multi-copy plasmids as their sole plasmid.

Therefore, plasmids with PCN > 10 mostly coexist with large low-copy plasmids (Binomial test: *p* < 10^−15^). Interactions among plasmids, in particular, interactions among small multi-copy plasmids and larger conjugative plasmids, may therefore play a role in the empirical scaling law between plasmid length and copy number.

### Genetic features associated with plasmid length and copy number

#### Length is more conserved than copy number within plasmid taxonomic groups (PTUs)

We assigned plasmids to PTUs using multiple, previously published plasmid classifications, to see how plasmid lengths and copy number vary with genetic distance. First, we used the PTUs reported by Acman et al.^42^ and Redondo-Salvo et al.^43^, who clustered plasmids into PTUs by *k*-mer similarity and Average Nucleotide Identity, respectively. In both cases, plasmids within a given PTU cluster by length, and have similar copy numbers (Supplementary Figure S4). Second, we clustered plasmids into PTUs based on similarity by Mash distance (using the default 0.06 threshold) using MOB-cluster^29^. Third, we typed plasmids based by their Rep proteins with MOB-typer^29^. Again, plasmids within PTUs based on these definitions have similar lengths and copy numbers. Finally, we used the Rep typing reported by Ares-Arroyo et al.^44^. Since plasmid incompatibility groups are largely defined by the Rep proteins that initiate plasmid replication, these results indicate that PCN strongly associates with plasmid length and the molecular systems that initiate plasmid replication. Across these plasmid classification systems, plasmid length is more conserved than PCN within PTUs. This finding indicates that highly related plasmids have similar lengths, but that the copy numbers for small plasmids can vary over an order of magnitude (Supplementary Figure S4).

#### Plasmid relaxase typing does not determine plasmid length and PCN

Plasmid lengths and copy numbers are not determined by plasmid mobility types as annotated by MOB-typer^29^, although some mobility groups are specific to large conjugative plasmids. Plasmid mobility groups are defined by the relaxase proteins that nick plasmids at *oriT* transfer origins to initiate horizontal gene transfer by conjugation. Specifically, many mobility groups show two modes, one corresponding to small mobilizable multi-copy plasmids, and a second corresponding to large low-copy conjugative plasmids (Supplementary Figure S5).

#### No correlations between PCN and plasmid host range

We also examined plasmid lengths and copy numbers in the context of host range annotation made by MOB-typer^29^ and the host range plasmid annotations reported by Redondo-Salvo et al.^43^ (Supplementary Figure S6). Large and small plasmids are found together across annotated host ranges, indicating that plasmid size and copy number does not correlate well with plasmid host range.

### Many plasmids have PCN < 1

Our data shows that many plasmids have a lower copy number than the chromosome (Figure 1, Supplementary Figure S7). Our methodological validations of PIRA (Supplementary Figure S1) show that this result is a genuine property of the sequencing data used for these PCN estimates. Out of the 11,338 plasmids in these data, 2,376 plasmids have PCN < 1, representing 21% of all plasmids. Across ecological categories, between 10% and 25% of plasmids have PCN < 1 (Supplementary Figure S7B).

Why do so many plasmids have PCN < 1? First, the PCN estimates reported in this work are relative to chromosome copy numbers, and do not represent estimates of the absolute number of plasmids per cell. It is known that bacteria can contain multiple chromosome copies per cell^45–47^, such that a clonal bacterial population can contain subpopulations with different numbers of chromosome copies per cell. The frequencies of these subpopulations can change over long-term evolution^48^.

Furthermore, some large bacteria show extreme polyploidy comprising tens of thousands of chromosome copies per cell^49^. This means that PCN estimates < 1 can arise in cases where say, a bacterium to has four chromosome copies, and two plasmid copies. In this scenario, we would estimate PCN = 0.5.

Second, PCN estimates < 1 could arise when genomic DNA is prepared from exponential phase cultures. During exponential growth, PCNs can be lower than chromosomal copy numbers^50,51^.

Third, this result could be caused by plasmid heterogeneity in sequenced bacterial clones, where some, but not all daughter cells contain the plasmid. In other words, the mixture of cells in an otherwise clonal sample may vary in the number of plasmids per cell, where the plasmid is present in some cells and absent from others. Such a scenario could arise during dynamic division of labor on plasmids, where a costly plasmid is maintained in a bacterial population by horizontal gene transfer from slow-growing producer cells to fast-growing cells without the plasmid^52^. PCN < 1 could also be maintained by the presence of parasitic satellite plasmids that effectively reduce the copy number of the plasmid of interest^53^.

### High copy number plasmids are rare and are enriched in human-impacted environments

High copy number plasmids (PCN > 50), by contrast, are relatively rare in this dataset (Supplementary Figures S7A and S7C). Out of the 11,338 plasmids in these data, 527 plasmids have PCN > 50, representing 4.6% of all plasmids. High copy number plasmids are significantly enriched in human-impacted environments (Supplementary Figure S7B). This finding suggests that very high PCNs (PCN > 50) may be transient phenomena generated by strong selection for the amplification of genes on small multicopy plasmids with stable copy numbers of ∼10-40 copies per chromosome per cell (Figure 1).

### Functional properties and organization of plasmids approach the functional organization of chromosomes as they increase in size

Our findings show how PCNs converge to chromosome copy numbers, as plasmids increase in length (Figure 1). We hypothesized that many functional properties of plasmids should converge to the functional properties of chromosomes as they increase in size. Indeed, we find that scaling laws emerge for the fraction of protein-coding sequences per plasmid and for the number of metabolic proteins per plasmid, as plasmids increase in length. These scaling laws converge to scaling laws that hold for chromosomes. This analysis was conducted on all complete microbial genomes containing plasmids in the NCBI RefSeq database, comprising 17,952 genomes containing 45,584 plasmids at the time of analysis.

#### Protein-coding sequence scaling law

As plasmids increase in size, the fraction of sequence dedicated to protein-coding sequences converges to the fraction of sequence dedicated to protein-coding sequences on chromosomes (Figure 2A and 2B). This pattern holds across plasmids and chromosomes sampled across diverse environments and microbial taxa across the tree of life (Figure 2C and Supplementary Figure S8). This finding implies that as plasmids increase in length, protein coding becomes more efficient. A simple explanation is that the overhead for regulatory sequences that determine plasmid replication, stability, and maintenance is relatively larger for small plasmids compared to large plasmids. For instance, a plasmid replication origin may take up some fixed length of DNA, and the relative length of this noncoding sequence decreases as a plasmid increases in length. In other words, as a plasmid increases in length, the relative length of the minimal sequence requirements for an autonomously replicating plasmid decreases, while this fraction may be relatively large for very small plasmids (Figure 2B).

**Figure 2.**
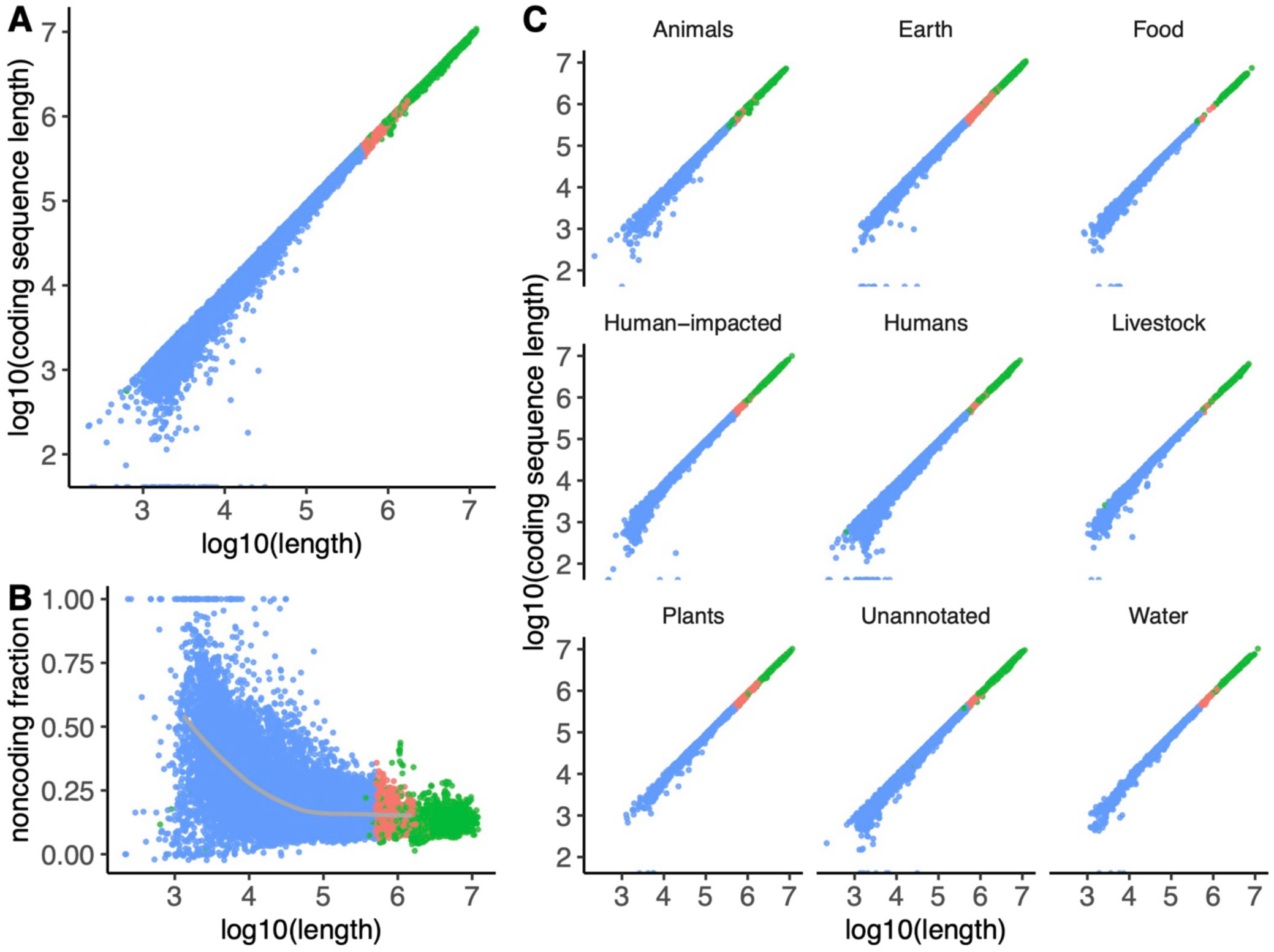
Protein-coding sequences on plasmids follow an empirical scaling law. Comparison of scaling between chromosomes and plasmids on a log-log scale. Plasmids are shown in blue, megaplasmids (plasmids > 500,000 bp in length) are shown in red, chromosomes are shown in green. **A)** As plasmids increase in size, the fraction of sequence dedicated to protein-coding sequences converges to the fraction of sequence dedicated to protein-coding sequences on chromosomes. **B)** As plasmids increase in size, the fraction of sequence dedicated to protein-coding sequences increases. Noncoding fraction is defined as (length – coding sequence length) / (length). The gray line indicates the average noncoding fractions and lengths of 100 buckets of plasmids, binned by length. **C)** The same pattern holds for microbes sampled across diverse environments. The ecological provenance of each replicon was annotated per the method described in Maddamsetti et al.3 (Methods).

#### Emergence of a metabolic scaling law as plasmids approach chromosome length scales

We annotated metabolic genes on plasmids by mapping genes to the metabolic pathways annotated in the KEGG database^54^, using the GhostKOALA functional annotation webserver^55^. Plasmid size scales with the number of metabolic genes on the plasmid. Given the computational cost of annotating chromosome metabolic genes with the GhostKOALA webserver, we only annotated metabolic genes for 100 chromosomes, arbitrarily chosen to span the full rank distribution of chromosomes by length. The metabolic scaling relationship found for these 100 chromosomes emerges among megaplasmids that are longer than 500,000 bp (Figure 3). Again, this emergent scaling law holds across plasmids and chromosomes sampled across diverse environments and microbial taxa across the tree of life (Figure 3B and Supplementary Figure S9). This scaling law may emerge due to fundamental constraints on cellular energetics. Megaplasmids may need more metabolic capacity (measured by the number of metabolic genes) to compensate for the metabolic burden required to maintain such large plasmids. Strikingly, the metabolic scaling law only fails for Mycoplasmatota (Supplementary Figure S9B); these bacteria are obligate pathogens that lack cell walls and include bacterial species with the smallest known cells, genomes, and metabolisms, even lacking key pathways like the TCA cycle^56^. This finding additionally supports our interpretation that the metabolic scaling law is caused by fundamental constraints on cellular energetics.

**Figure 3.**
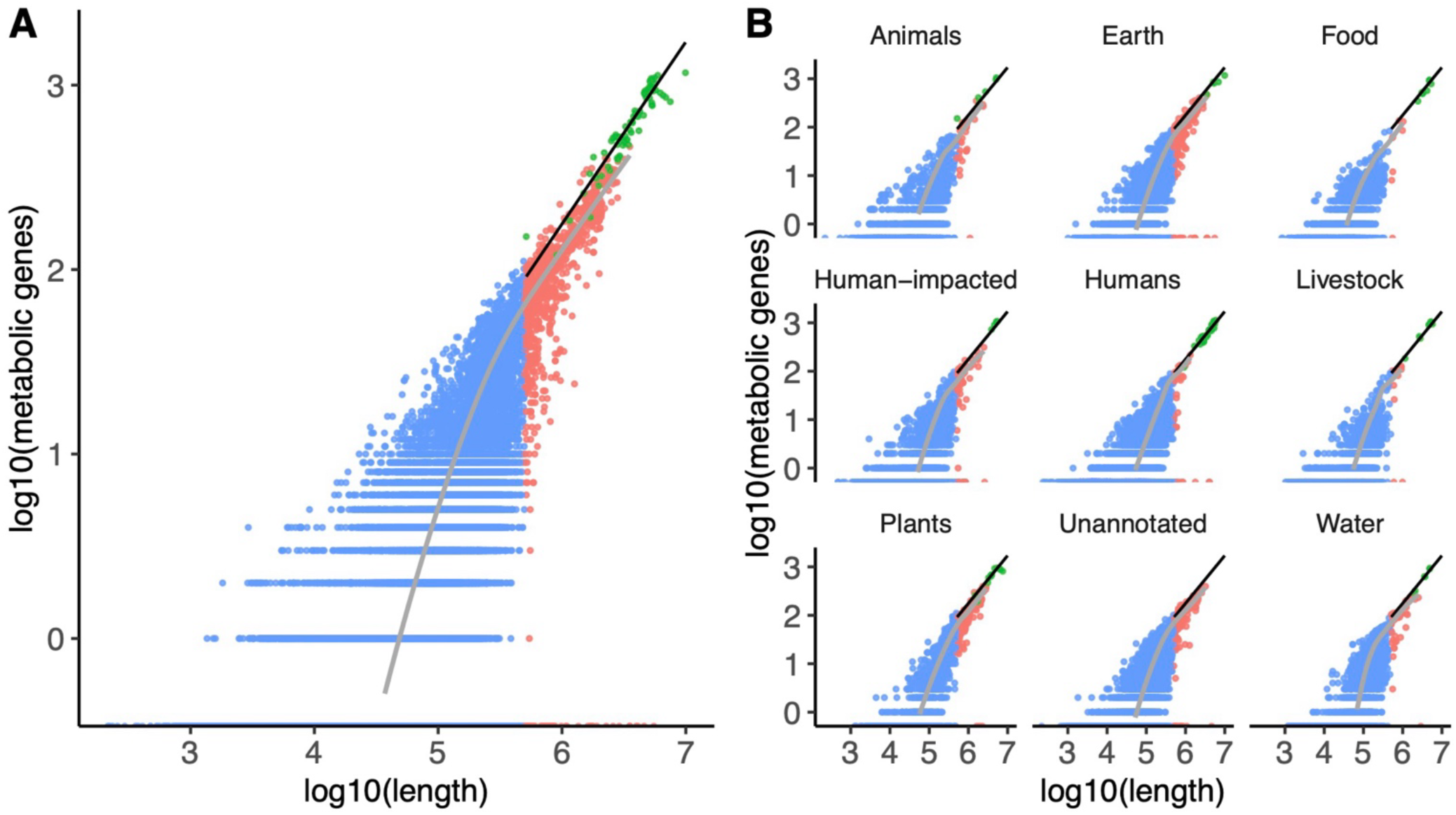
A metabolic scaling law emerges as plasmids approach chromosome length scales. Comparison of scaling between chromosomes and plasmids on a log-log plot. Plasmids are shown in blue, megaplasmids (plasmids > 500,000 bp in length) are shown in red, chromosomes are shown in green. The linear regression between log10(metabolic genes) and log10(length) for chromosomes is shown in black. The gray line indicates the average length of plasmids containing *n* metabolic genes, where *n* is an arbitrary integer. As *n* increases, the scaling between plasmid length and the number of metabolic genes *n* (shown in gray) approaches the scaling for chromosome length and number of metabolic genes *n* (shown in black). **A)** Plasmids can vary in size by orders of magnitude but still carry similar numbers of metabolic genes— but as plasmids reach megaplasmid scales (> 500,000 bp in length), their metabolic gene content begins to scale like chromosomes. **B)** The emergent metabolic scaling law holds for microbes sampled across diverse environments. The ecological provenance of each replicon was annotated per the method described in Maddamsetti et al.^3^ (Methods).

## Discussion

We have uncovered universal scaling laws that govern fundamental aspects of plasmid biology, revealing deep-rooted principles underlying microbial evolution and genome organization. This work reports the largest dataset on PCN and length to date, encompassing 4,317 bacterial and archaeal genomes and 11,338 plasmids. Our discovery of an inverse power-law correlation between PCN and plasmid length indicates a fundamental constraint that dictates how much plasmid DNA can be stably transmitted over time: once plasmid DNA content reaches ∼2% per chromosome, plasmid replication needs to be synchronized with the cell cycle for stable inheritance. This scaling law not only advances our understanding of plasmid biology but also challenges traditional distinctions between plasmids and chromosomes, suggesting a continuum of genetic elements shaped by fundamental biophysical constraints.

While we were finalizing our manuscript, we became aware of independent work reporting both a large dataset of PCNs as well as a scaling law relating PCN to plasmid length^57^. The convergence of findings from independent studies validates the universality of the inverse scaling law between plasmid length and copy number.

Our analyses also reveal scaling laws constraining fundamental aspects of plasmid biology, such as protein-coding regions and metabolism. We found that as plasmids increase in size, their functional properties and organization converge toward those of chromosomes, highlighting a unifying principle in genome evolution. These findings are consistent with previous reports that the functional gene content of microbial genomes follow empirical scaling laws^58–63^, and show that such relations extend to plasmids. The emergent metabolic scaling law, in particular, indicates that larger plasmids incorporate more metabolic genes, suggesting that metabolic capacity is a critical factor in the stable maintenance of large genetic elements. Together, our results demonstrate the existence of fundamental constraints that dictate stable plasmid coding structures. These constraints have broad implications for the evolution of microbial genomes, the dynamics of horizontal gene transfer, and the adaptability of microbial communities in various environments.

Moreover, our findings have practical implications for biotechnology and synthetic biology. Understanding the scaling laws governing plasmid biology can inform the rational design of synthetic plasmids, optimizing them for desired functions while ensuring stable inheritance and minimal metabolic burden on host cells^11,28^. One intriguing implication of the scaling laws we uncovered is that larger plasmids may serve a more efficient chassis for engineered functions. As plasmid size increases, the fraction of DNA dedicated to protein-coding sequences and metabolic genes scales predictably, converging toward chromosomal characteristics. This suggests that larger plasmids have greater potential to accommodate diverse functional modules, particularly those related to metabolic processes. By contrast, smaller plasmids appear to require a higher fraction of their noncoding DNA for essential functions such as replication, stability, and maintenance (Figure 2B).

This observation could indicate that small plasmids are more constrained in their ability to host engineered gene circuits, as the available space for protein-coding sequences is more limited.

This scaling behavior raises the possibility that larger plasmids are inherently more efficient for applications where high encoding capacity is required. Not only can they support more protein-coding sequences, but they may also be more suited to carry complex metabolic pathways. By using larger plasmids as chassis, synthetic biologists might achieve more stable and scalable designs with reduced competition between essential functions and engineered traits. This finding could help guide the selection of plasmid sizes in the design of synthetic constructs, favoring larger plasmids for applications requiring extensive metabolic or functional gene integration^64,65^.

Our work also demonstrates the power of applying efficient algorithms (e.g., pseudoalignment) developed for one problem (transcriptomics), to another (PCN estimation) to reveal new biology. For example, pseudoalignment could accelerate the inference of microbial growth rates from peak-to-trough coverage ratios in microbiome data^66^, as well as the discovery of structural variation in microbiomes^67^, as the state-of-the-art currently depends on alignment-based methods^66,67^. Based on our current results, we hypothesize that such an analysis would show that plasmid replication rates converge to chromosome replication rates in microbiome data, as plasmid lengths approach the size of chromosomes. Second, pseudoalignment should accelerate the discovery of high-copy number plasmids and gene amplifications in large databases of metagenome-assembled genomes (MAGs)^68–70^ . By rapidly remapping raw sequencing data to MAGs using PIRA or other pseudoalignment-based methods, the relative copy numbers of different replicons in a metagenomic sample may be estimated. This approach could facilitate the identification of genetic elements associated with antibiotic resistance, virulence factors, or metabolic capabilities, with significant implications for public health and environmental microbiology. The discovery of replicons with elevated copy number in microbiome data may represent high-copy-number plasmids, mobile genetic elements, viruses, or genetic amplifications mediated by selection and horizontal gene transfer^2,3^.

In summary, our study uncovers universal scaling laws that govern plasmid biology, revealing fundamental constraints on plasmid evolution and function. By developing and applying PIRA, we have not only expanded our understanding of microbial genetics but also provided tools and insights that can drive future research and applications in biotechnology, medicine, and environmental science. Our findings emphasize the interconnectedness of genetic elements and highlight the importance of considering fundamental biophysical constraints in the study of genome evolution.

## Materials and Methods

### Plasmid copy number analysis

#### Input genomes for the plasmid copy number analysis pipeline

A table of microbial genomes in the NCBI genomes database was downloaded from: https://ftp.ncbi.nlm.nih.gov/genomes/GENOME_REPORTS/prokaryotes.txt. This table was filtered for genomes containing plasmids by running the following UNIX command: *(head -n 1 ../data/prokaryotes.txt && grep "plasmid" ../data/prokaryotes.txt | grep "chromosome") > ../results/prokaryotes-with-chromosomes-and-plasmids.txt*. This command ensures that every genome has an annotated chromosome and at least one annotated plasmid. This resulted in a filtered table of 19,538 microbial genomes containing plasmids. These genomes were then filtered by the availability of corresponding Illumina short-read sequence data in the NCBI Sequencing Read Archive (SRA).

#### Downloading of reference genome annotation and corresponding Illumina short-read sequencing data

Data downloading was automated in the Python 3.11 script *PCN_pipeline.py*. First, metadata for each genome in *prokaryotes-with-chromosomes-and-plasmids.txt* was examined to find the subset of genomes with paired-end Illumina sequencing data in the NCBI Sequencing Read Archive.

This resulted in a table, *RunID_table.csv*, containing records for 4,921 genomes containing plasmids. The NCBI RefSeq database was cross-checked against these genomes, and reference genome annotation data was downloaded for the subset of microbial genomes found in the NCBI RefSeq database. Illumina paired-end short read sequencing data in *fastq* format was downloaded for each of these microbial genomes in RefSeq; this step was the most time-consuming step of this pipeline, taking two weeks to download ∼15 TB of sequencing data, using the *prefetch* and *fasterq-dump* programs in NCBI SRA Toolkit v3.0.5^20^. At this stage, a Python 3.11 script called *check- genome-quality-and-consistency.py* was used to ensure that all downloaded *fastq* data corresponded to a downloaded reference genome, and to ensure that all reference genomes contained complete chromosome assemblies. In all, data for 4,540 microbial genomes in NCBI RefSeq were downloaded.

#### Pseudoalignment of sequencing reads against reference genomes and direct PCN estimation

Sequencing data processing was automated in the Python 3.11 script *PCN_pipeline.py*. *kallisto 0.48.0* and *themisto 3.2.2* were used to pseudoalign Illumina sequencing reads against reference chromosomes and plasmids. Sequencing coverage per replicon (i.e. chromosome or plasmid) was estimated by dividing the number of reads mapping to the replicon by the length of replicon. Then, direct estimates of PCNs (relative to chromosome) were generated by dividing the mean sequencing coverage for each plasmid by the mean sequencing coverage for the largest chromosome in each genome.

#### Probabilistic Iterative Read Assignment (PIRA)

PIRA was developed to further increase the accuracy of PCN estimates by incorporating information from sequencing reads that map ambiguously to multiple replicons within a genome, such as a read that maps to both a plasmid and chromosome. This situation can arise when reads come from repeated or duplicated sequences, as in the case of a read that corresponds to a transposon found in multiple locations in a genome. A specification of the PIRA algorithm is provided in Supplementary Data 1, and PIRA is implemented in the Python 3.11 script *PCN_pipeline.py*. For a given genome, an initial copy number estimate vector is generated using the direct method described in the previous section. This initial copy number estimate vector ignores multireads that pseudoaligned to multiple replicons. The original *fastq* files are then filtered for multireads, and these multireads are re-aligned to the reference genome using *minimap2* version 2.28-r1209^24^. Reads that align to a single location by traditional sequencing alignment with *minimap2* are tabulated. These are used to further improve the initial copy number estimate vector. The remaining *m* multireads are then tabulated into an *m* × *n* matrix, where *n* is the number of replicons in the genome, sorted by length, such that the longest replicon (that is, the main chromosome) is in the first position. Each row of the matrix (each row corresponding to a single multiread) has entries corresponding to the number of times that this multiread maps to each replicon in the genome.

The columns of this matrix are scaled by the entries of the initial copy number vector, so that each row (corresponding to a single multiread) accounts for both the number of locations on the replicon that align with this multiread, as well as the current copy number estimate of each replicon. Each row is then normalized to sum to one, such that each row now represents the probability distribution of which replicon that multiread came from. Then, all rows are summed to make a 1 × *n* vector that sums to *m*; this vector represents how all the multireads probabilistically map to the replicons. This vector of multiread counts is used to update the copy number estimate vector (by adding these multiread counts to the number of unireads mapping to the replicon, and dividing the total by replicon length), and the process is iterated until the copy number estimate vector converges (i.e., the norm of the change between iterations falls below a small value like 10^−6^).

### Comparison with the ICRA algorithm

At a high-level, PIRA is in the same class of methods as the “iterative coverage-based read assignment” (ICRA) algorithm described by Zeevi et al.^67^, but differing in four aspects. First, PIRA was designed to infer copy numbers within genomes, and not within metagenomes. Second, PIRA lets a multiread contribute to multiple replicons based on the probability that the read originated from that replicon, rather than assigning a multiread to the single best match (“soft” read assignment, rather than “hard” read assignment, in machine learning parlance). Third, PIRA weights the probability distribution of how a multiread maps to replicons within a genome by 1) the number of matches of that read to a given replicon and 2) the current estimate of that replicon’s copy number; unlike ICRA, alignment read quality is not considered.

Fourth, ICRA divides replicons into genomic bins, and iteratively estimates the copy numbers of each genomic bin to find copy number variation; by contrast, PIRA treats each replicon as a single bin. In principle, PIRA could be generalized to estimate copy number within genomic bins like ICRA: this is a direction for future research. While PIRA was conceived independently from ICRA, its similarities to ICRA gave us confidence that PIRA was the correct approach for incorporating multiread information. In addition, the two-dimensional matrix data structure used to store multireads for PIRA was conceived as a simpler version of the 3-dimensional array described by Zeevi et al. in their formal description of ICRA^67^.

#### Benchmarking of PCN estimates

We benchmarked our PCN estimate pipeline using a random subset of 100 genomes, each containing at least one plasmid with an estimated PCN < 0.8. This benchmarking was automated in the Python 3.11 script *PCN_pipeline.py*. First, PCNs were re-estimated on these 100 genomes using *minimap2* version 2.28-r1209 to align all reads, rather than using pseudoalignment. PIRA was used to account for multireads found with *minimap2*. Second, PCNs were re-estimated on these 100 genomes using *breseq* 0.35 to align all reads. In this case, multireads were ignored, and PCNs were estimated by dividing plasmid mean sequencing coverage by chromosome mean sequencing coverage. We also compared PCN estimates with the direct method (ignoring multireads) using *themisto* 3.2.2 to pseudoalign reads, and PCN estimates using PIRA on top of the direct method using *themisto*. We also compared direct PCN estimates using *Themisto* 3.2.2 with direct PCN estimates using *kallisto* 0.48.0 to see how the choice of pseudoalignment software affected PCN estimates, if at all.

### Ecological annotation

The ecological provenance of each microbial genome was annotated per the method described in Maddamsetti et al.^3^. Briefly, we used the “host” and “isolation_source” fields in the RefSeq annotation for each genome to place each into the following categories: Marine, Freshwater, Human-impacted (environments), Livestock (domesticated animals), Agriculture (domesticated plants), Food, Humans, Plants, Animals (non-domesticated animals, also including invertebrates, fungi and single-cell eukaryotes), Soil, Sediment (including mud), Terrestrial (non-soil, non-sediment, including environments with extreme salinity, aridity, acidity, or alkalinity), and Unannotated (no annotation). For reproducibility, our annotations are generated using a Python 3.11 script called *annotate-ecological-category.py* and checked for internal consistency using a Python 3.11 script called *check-ecological-annotation.py*. To simplify the data presentation, we merged categories as follows. Marine and Freshwater categories were grouped as “Water”. Sediment, Soil, and Terrestrial categories were grouped as “Earth”. Plant and Agriculture categories were grouped as “Plants”.

### Estimation of PCN reporting in the scientific literature

A random seed was drawn between 1 and 100 using the random number generator at www.random.org^71^. For replicability, this random seed (the number 60) was used to draw 50 genomes without replacement, from a set of 1216 genomes containing at least one plasmid with an estimated PCN greater than 10. The RefSeq annotation files for each of these genomes was examined manually for publications associated with the given genome assembly. Each publication was examined manually to see if PCNs were reported. The results of this analysis are reported in Supplementary Table S1.

### Plasmid scaling law analysis

#### Input genomes for the plasmid scaling law analysis

A table of microbial genomes in the NCBI genomes database was downloaded from: https://ftp.ncbi.nlm.nih.gov/genomes/GENOME_REPORTS/prokaryotes.txt. This table was filtered for complete microbial genomes containing plasmids by running the Python 3.11 script *filter- genome-reports.py*. This resulted in a filtered table of 17,952 complete microbial genomes containing 45,584 plasmids. Reference genome annotation files for these genomes were downloaded using the Python 3.11 script *fetch-gbk-annotation.py*. Then, the following python 3.11 scripts were run to tabulate ecological metadata, and protein counts and lengths for each genome: *make-chromosome-plasmid-table.py*, *make-gbk-annotation-table.py*, and *count-proteins-and- replicon-lengths.py*.

#### Protein-coding sequence calculations

The fraction of protein-coding sequences per replicon was calculated by summing up the length of each protein in each replicon (i.e., plasmid or chromosome) in each genome, and dividing by the length of that replicon. This calculation was run using the Python 3.11 script *calculate-CDS-rRNA-fractions.py*.

#### Analysis of metabolic genes on plasmids

We annotated metabolic genes on plasmids following the computational protocol in Hamrick et al.^52^. Briefly, the GhostKOALA functional genomics web server^55^ (https://www.kegg.jp/ghostkoala/) associated with the Kyoto Encyclopedia of Genes and Genomes (KEGG) database^54^ was used to annotate plasmid proteins with KEGG Orthology (KO) IDs. The Python 3.11 scripts *make-plasmid-protein-FASTA-db.py* and *make-plasmid-GhostKOALA- input-files.py* were used to generate input files for GhostKOALA. Each of the these input files were manually submitted to the GhostKOALA webserver at https://www.kegg.jp/ghostkoala, and saved to disk. Then, the shell script *concatenate-and-filter-plasmid-GhostKOALA-results.sh* was used to concatenate the corresponding GhostKOALA output files and filter for plasmid proteins that were successfully mapped to a KEGG KO ID. The Python 3.11 script *remove-chromosomes-from- plasmid-Ghost-KOALA-results.py* was used to remove any proteins found on plasmids larger than chromosomes, as these were assumed to be annotation errors. The, the union of all KEGG KO IDs found among the plasmid genes was generated using the Python 3.11 script *get_unique_KEGG_IDs.py*. The output of this script was uploaded to the KEGG Mapper Reconstruct web server at: https://www.kegg.jp/kegg/mapper/reconstruct.html. The set of plasmid KEGG KO IDs that mapped to metabolic pathways (KEGG PATHWAY Database ID: 01100) was saved to file. Then, the Python 3.11 script *get-plasmid-metabolic-KOs.py* was run to generate a table of all metabolic genes on plasmids.

#### Analysis of metabolic genes on chromosomes

For comparison to the analysis of metabolic genes on plasmids, we examined metabolic genes on the chromosomes of 100 representative and arbitrarily chosen genomes. Genomes were ranked based on the length of their chromosome, and 100 genomes were chosen, by virtue of being roughly equally distributed across the rank-distribution of chromosome lengths, over the set of genomes with ecological annotation (i.e., genomes marked as “Unannotated” were never chosen), and the proteins found in these 100 genomes were put into a small database file using the Python 3.11 script *make-chromosome-protein-FASTA-db.py*. This input file was submitted to the GhostKOALA web server, and the GhostKOALA output file was saved to disk. Then, the shell script *filter-chromosome-GhostKOALA-results.sh* was used to filter the GhostKOALA output for chromosomal proteins that were successfully mapped to a KEGG KO ID. The Python 3.11 script *get_unique_chromosome_KEGG_IDs.py* was used to get the union of all KEGG IDs found among these chromosomal proteins. These data were uploaded to the KEGG Mapper Reconstruct webserver (https://www.kegg.jp/kegg/mapper/reconstruct.html), and the set of chromosomal KEGG KO IDs that mapped to metabolic pathways (KEGG PATHWAY Database ID: 01100) was saved to file. Then, the Python 3.11 script *get-chromosome-metabolic-KOs.py* was used to generate a final table of all KEGG metabolic proteins in the sampled chromosomes.

### Plasmid typing metadata

All plasmids were annotated using *MOB-typer 3.1.7*^29^ using the protocol reported by Hamrick et al.^52^. The annotations used in this work were previously reported in Supplementary Data File 5 of Maddamsetti et al.^3^. Plasmids were also typed using the supplementary data reported by Acman et al.^42^, Redondo-Salvo et al.^43^, Coluzzi et al.^72^, and Ares-Arroyo et al.^44^.

### Statistical analysis

All statistical analysis and data visualizations were generated using an R 4.2 script called *PCN- analysis.R*. At this stage, plasmid sequences < 1000 bp in length were removed from the analysis to remove unmapped or unplaced plasmid contigs. Genomes with plasmids longer than their chromosome were also removed, to remove potential genome misannotation errors.

### Data Availability

All data analyzed in this project was retrieved from the NCBI RefSeq and Sequencing Read Archive databases, and is therefore publicly available.

### Code Availability

A Github repository containing all data and code sufficient to reproduce the PCN estimation pipeline, including downloading of genomic data, is available at: https://github.com/rohanmaddamsetti/PCN-db-pipeline. A Github repository containing all computer codes for the remaining analyses, including statistics and figures, is available at: https://github.com/rohanmaddamsetti/plasmid-scaling-laws.

## Acknowledgements

We thank the members of the You lab for helpful discussions and comments, and Duke Research Computing for technical assistance and computing resources. This work is partially supported by the National Institutes of Health (L.Y., R01AI125604, R01GM098642, and R01EB031869). The funders had no role in study design, data collection and analysis, decision to publish, or preparation of the manuscript.

## Author Contributions

RM and LY conceived and designed research. RM, MLW, HS, and ZZ conducted bioinformatic analyses. JL checked PIRA for correctness. RM and LY wrote the manuscript. RM, JL, ZZ, and LY edited the manuscript. LY supervised research.

## SUPPLEMENTARY INFORMATION

**Supplementary Data 1: PDF file of PIRA algorithm specification.**

**Supplementary Data 2: CSV file of plasmid copy number estimates.**

**Supplementary Figure S1.**
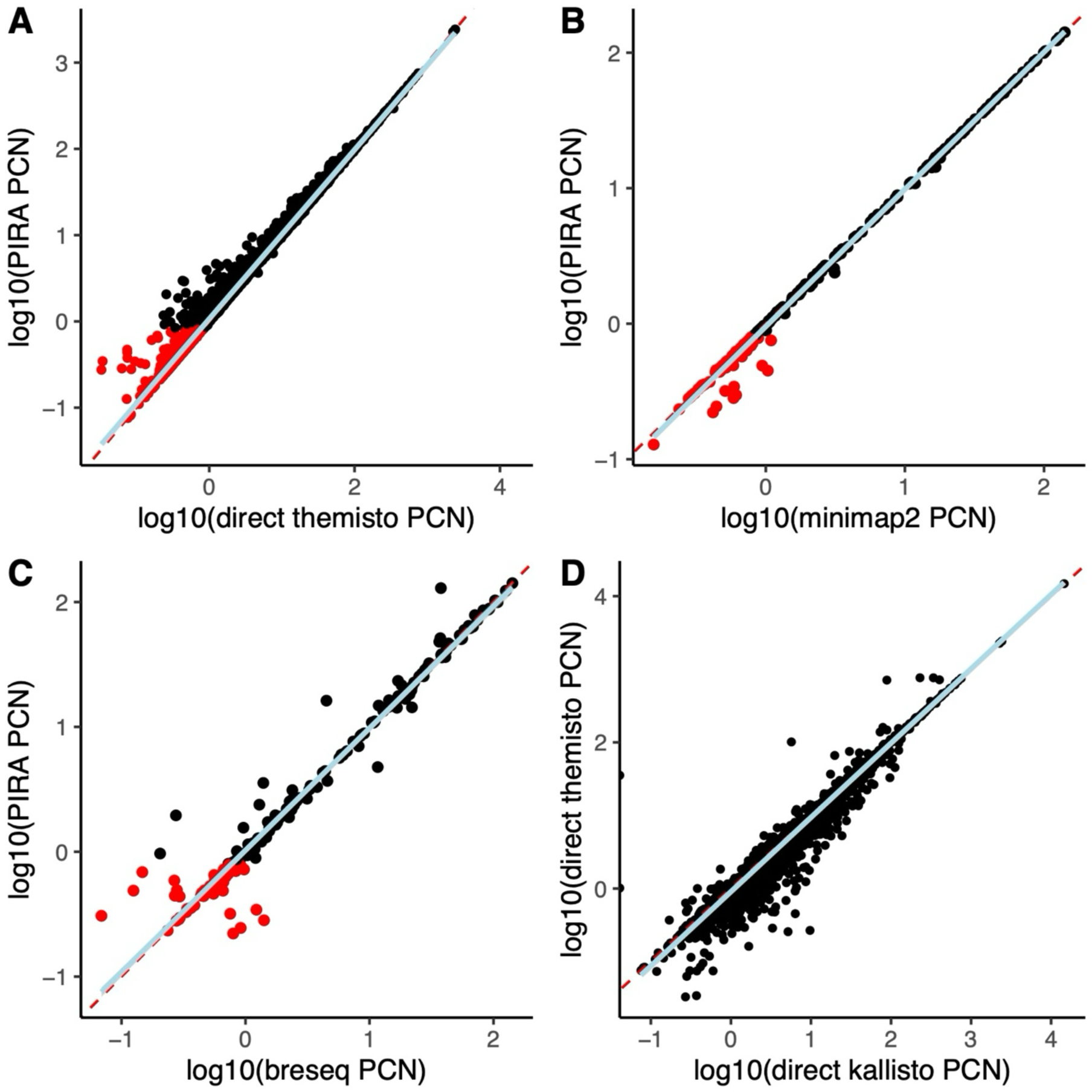
Plasmid copy number estimation benchmarking. Plasmids with PIRA PCN < 0.8 are labeled in red. In each panel, a linear regression showing the observed line of best fit is drawn in light blue. A dashed red line indicates the expected one-to-one correlation. These results indicate that PIRA provides accurate and reliable PCN estimates and show that the low PCN estimates (PCN < 0.8) are a property of the underlying sequencing data, given the consistency of these estimates across methods. **A)** Comparison of direct Themisto PCN estimates to PIRA PCN estimates, excluding plasmids with fewer than 10,000 mapped reads by the naïve method. **B)** Benchmarking of PIRA against minimap2. PCN estimates were compared between PIRA and minimap2 on a test set of 100 randomly selected genomes, each with at least one plasmid with PIRA PCN < 0.8. **C)** Benchmarking of PIRA against *breseq*. PCN estimates were compared between PIRA and *breseq* on a test set of 100 randomly selected genomes, each with at least one plasmid with PIRA PCN < 0.8. **D)** Comparison of direct *themisto* PCN estimates to direct *kallisto* PCN estimates.

**Supplementary Figure S2.**
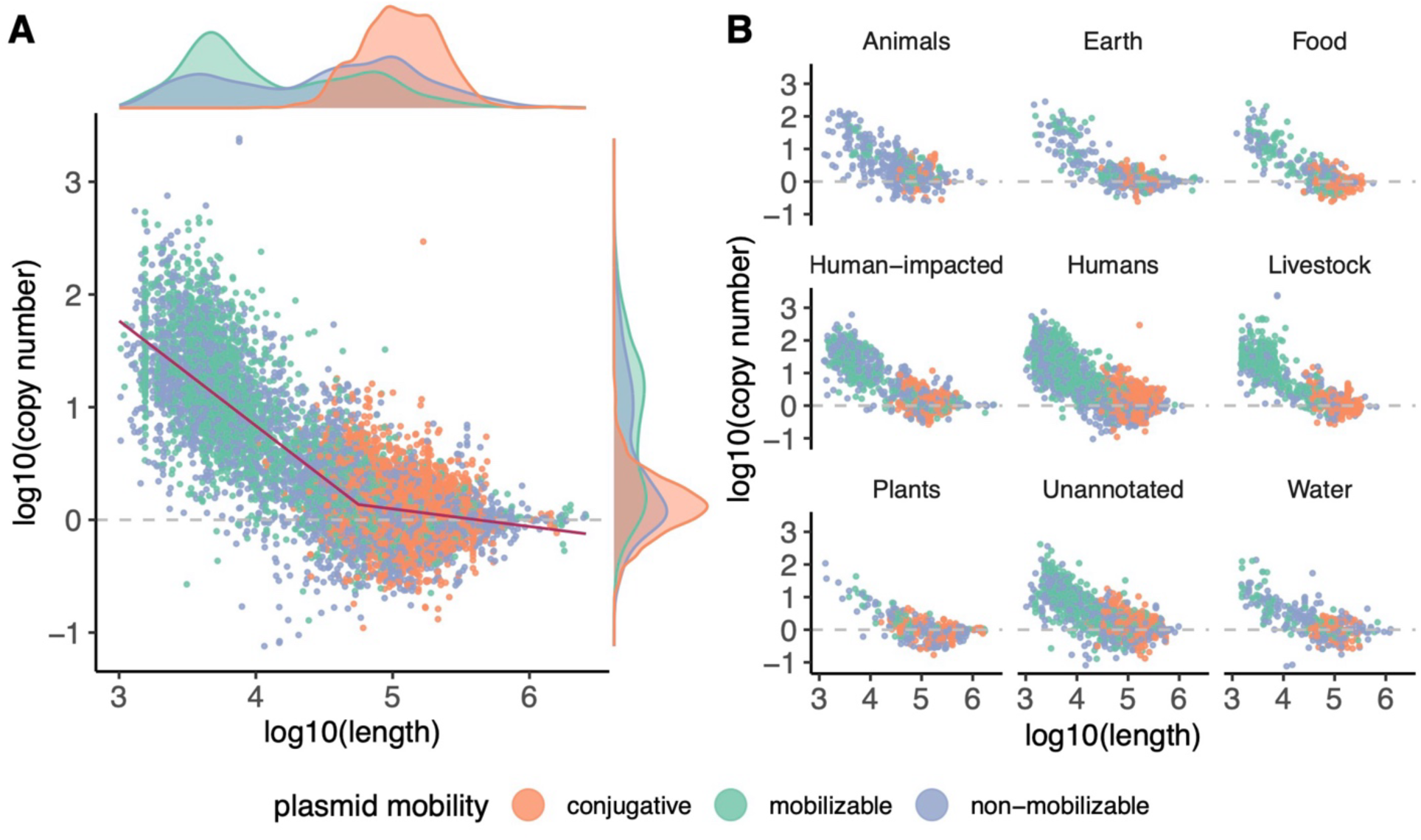
Plasmid length inversely correlates with plasmid copy number. **A)** Plasmid length inversely correlates with plasmid copy number. Even without rescaling plasmid length by the length of the largest chromosome, as in Figure 1, a biphasic scaling law is apparent. A segmented regression (in maroon) was fit to these data on a log-log plot. This segmented regression has a first slope of –0.93, a breakpoint at 4.56, a second slope of –0.15, and an Adjusted R^2^ of 0.664. The marginal density distributions of plasmid copy number and length are displayed on the axes. **B)** The inverse correlation between plasmid length and copy number holds across diverse environments.

**Supplementary Figure S3.**
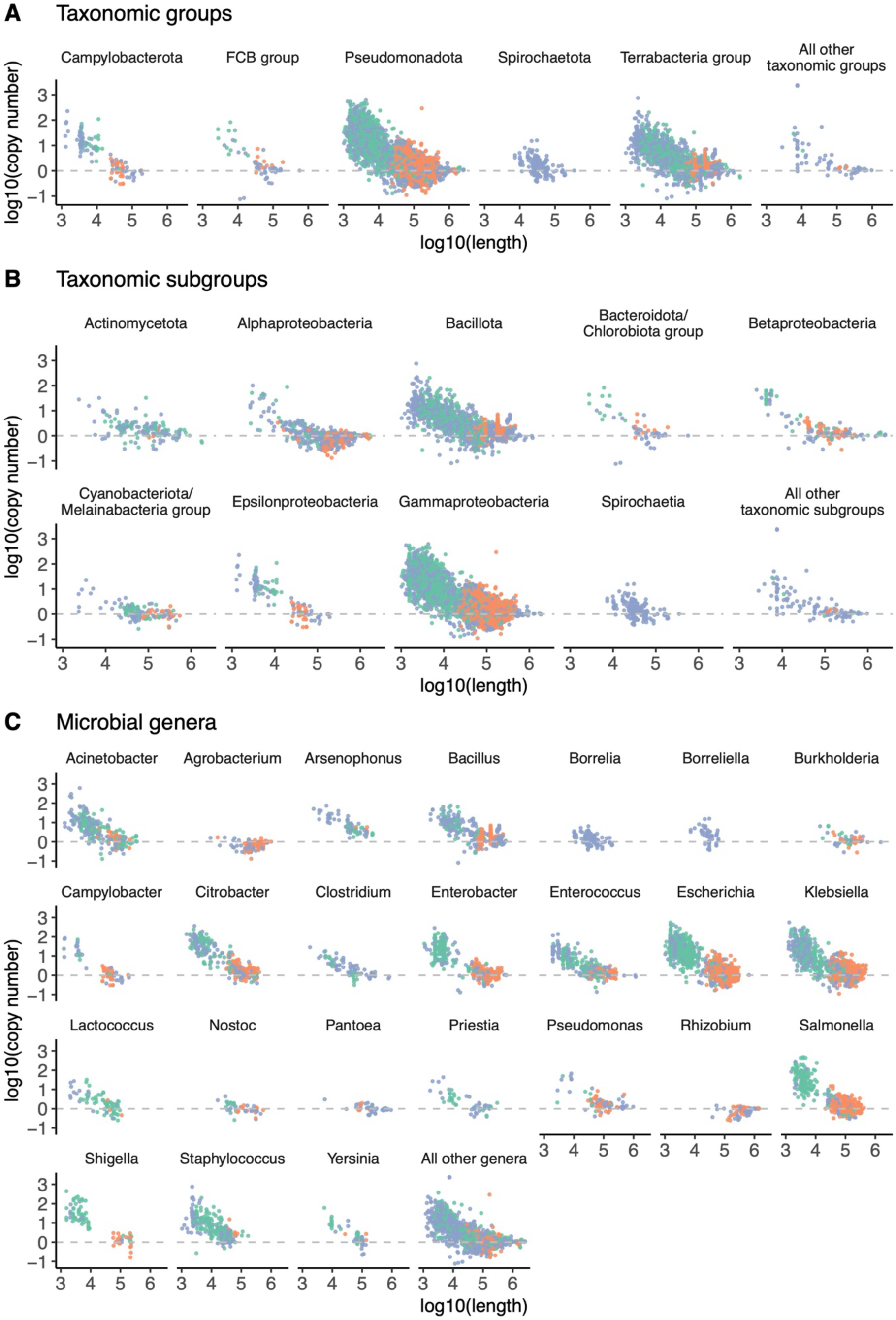
The inverse correlation between plasmid copy number and plasmid length holds across genera. Conjugative plasmids are colored orange pink, mobilizable plasmids are colored light green, and non-mobilizable plasmids are colored light blue. **A)** Each panel represents a NCBI taxonomic group with at least 50 plasmids; genera with fewer than 50 plasmids are lumped together in the final panel. **B)** Each panel represents a NCBI taxonomic subgroup with at least 50 plasmids; genera with fewer than 50 plasmids are lumped together in the final panel. **C)** Each panel represents a genus with at least 50 plasmids; genera with fewer than 50 plasmids are lumped together in the final panel.

**Supplementary Figure S4.**
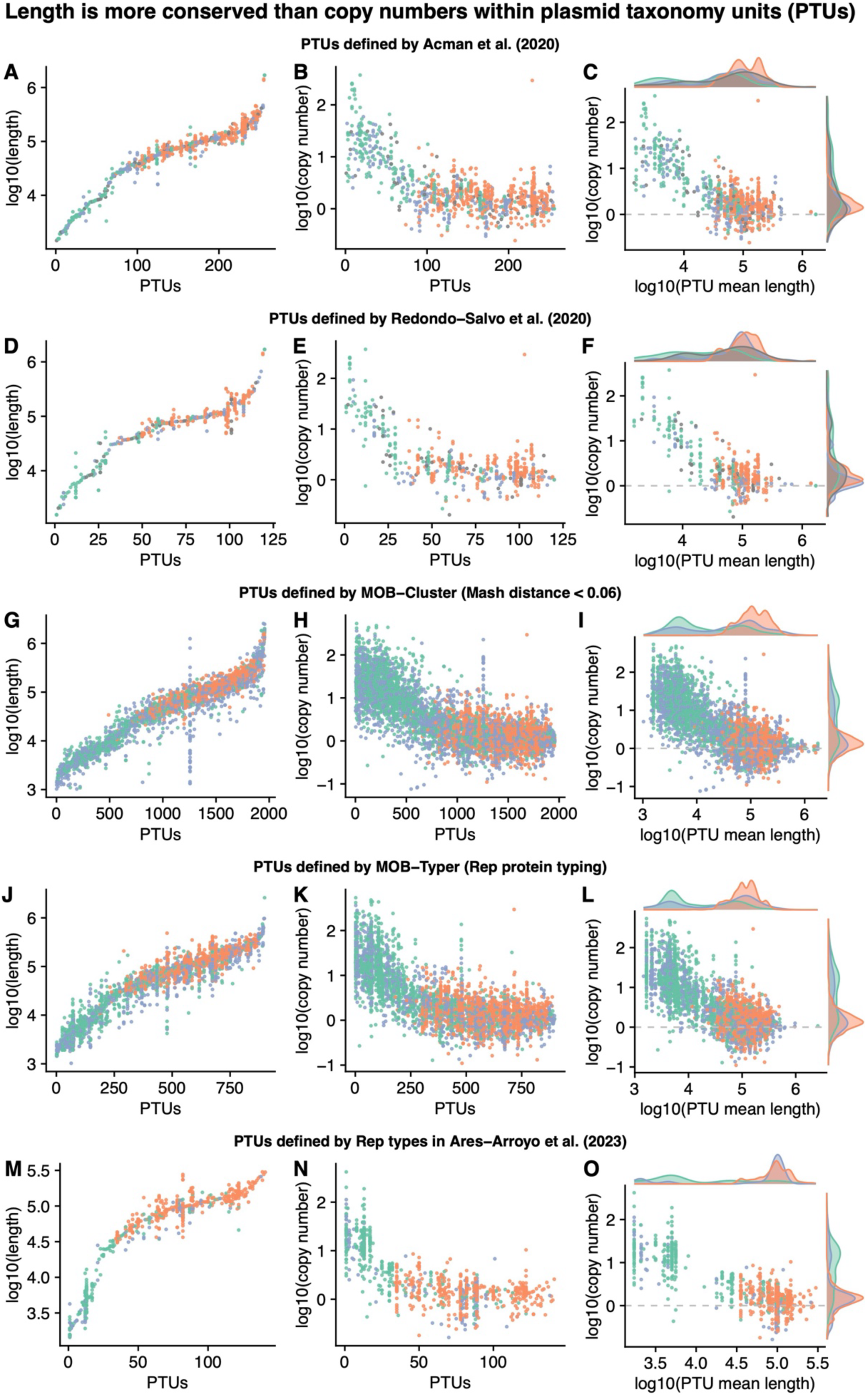
Length is more conserved than copy number within plasmid taxonomic groups (PTUs). Conjugative plasmids are colored orange pink, mobilizable plasmids are colored light green, and non-mobilizable plasmids are colored light blue. Each row represents a plasmid classification scheme described by different groups, to show the robustness of this finding to the choice of classification method. In every panel, PTUs are ranked by length from smallest to largest, to show variation within PTUs. The left panel shows variation in plasmid lengths within PTUs (i.e., per each rank on the x-axis). The right panel shows variation in plasmid copy numbers within PTUs (i.e., per each rank on the x-axis). **A)** Plasmid taxonomic units (PTUs) defined by the similarity network analysis by Acman, et al. (2020) cluster by length. **B)** PTUs defined by the similarity network analysis by Acman, et al. (2020) vary more in copy number than length. **C)** The inverse correlation between length and copy number holds across PTUs defined by Acman, et al. (2020). **D)** PTUs defined by the similarity network analysis by Redondo-Salvo, et al. (2020) cluster by length. **E)** PTUs defined by the similarity network analysis by Redondo-Salvo, et al. (2020) vary more in copy number than length. **F)** The inverse correlation between length and copy number holds across PTUs defined by Redondo-Salvo, et al. (2020). **G)** PTUs defined by MOB-Cluster cluster by length. **H)** PTUs defined by MOB-Cluster vary more in copy number than length. **I)** The inverse correlation between length and copy number holds across PTUs defined by MOB-Cluster. **J)** PTUs defined by MOB-Typer Rep protein typing cluster by length. **K)** PTUs defined by MOB-Typer Rep protein typing vary more in copy number than length. **L)** The inverse correlation between length and copy number holds across PTUs defined by MOB-Typer. **M)** PTUs defined by the Rep protein typing in Ares-Arroyo, et al. (2023) cluster by length. **N)** PTUs defined by the Rep protein typing in Ares-Arroyo, et al. (2023) vary more in copy number than length. **O)** The inverse correlation between length and copy number holds across PTUs defined by Ares- Arroyo, et al. (2023).

**Supplementary Figure S5.**
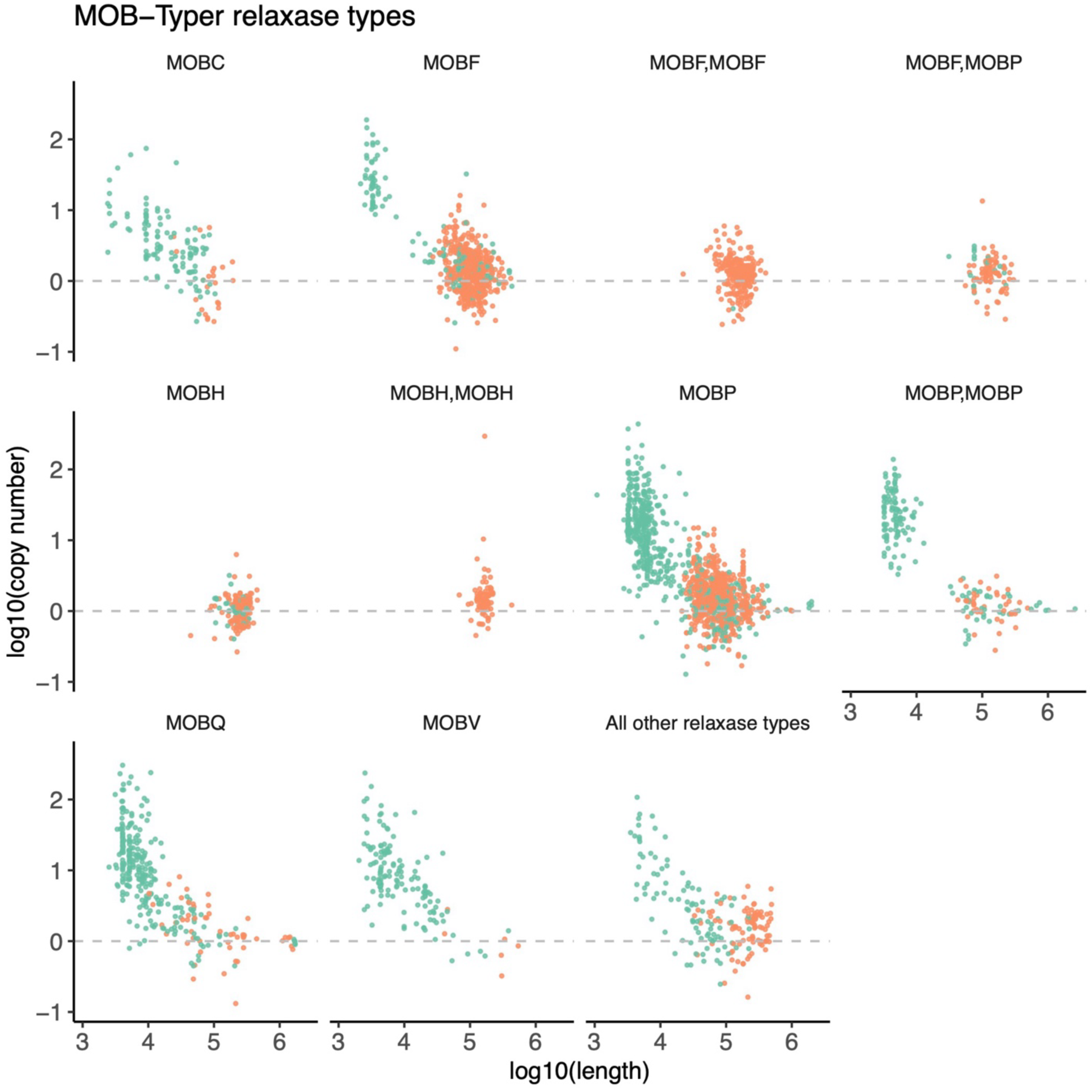
Plasmid mobility groups defined by relaxase types with MOB-Typer often contain both small mobilizable plasmids and large conjugative plasmids. Conjugative plasmids are colored orange pink and mobilizable plasmids are colored light green. Relaxase types with more than 50 plasmids are shown in separate panels; the remaining relaxase types are lumped together in the final panel. Some plasmids have multiple relaxase enzymes and may therefore belong to multiple mobility groups. For instance, the MOBF,MOBF panel indicates plasmids that contain two MOBF-family relaxases, and the MOBF,MOBP panel indicates plasmids that contain one MOBF-family relaxase and one MOBP-family relaxase.

**Supplementary Figure S6.**
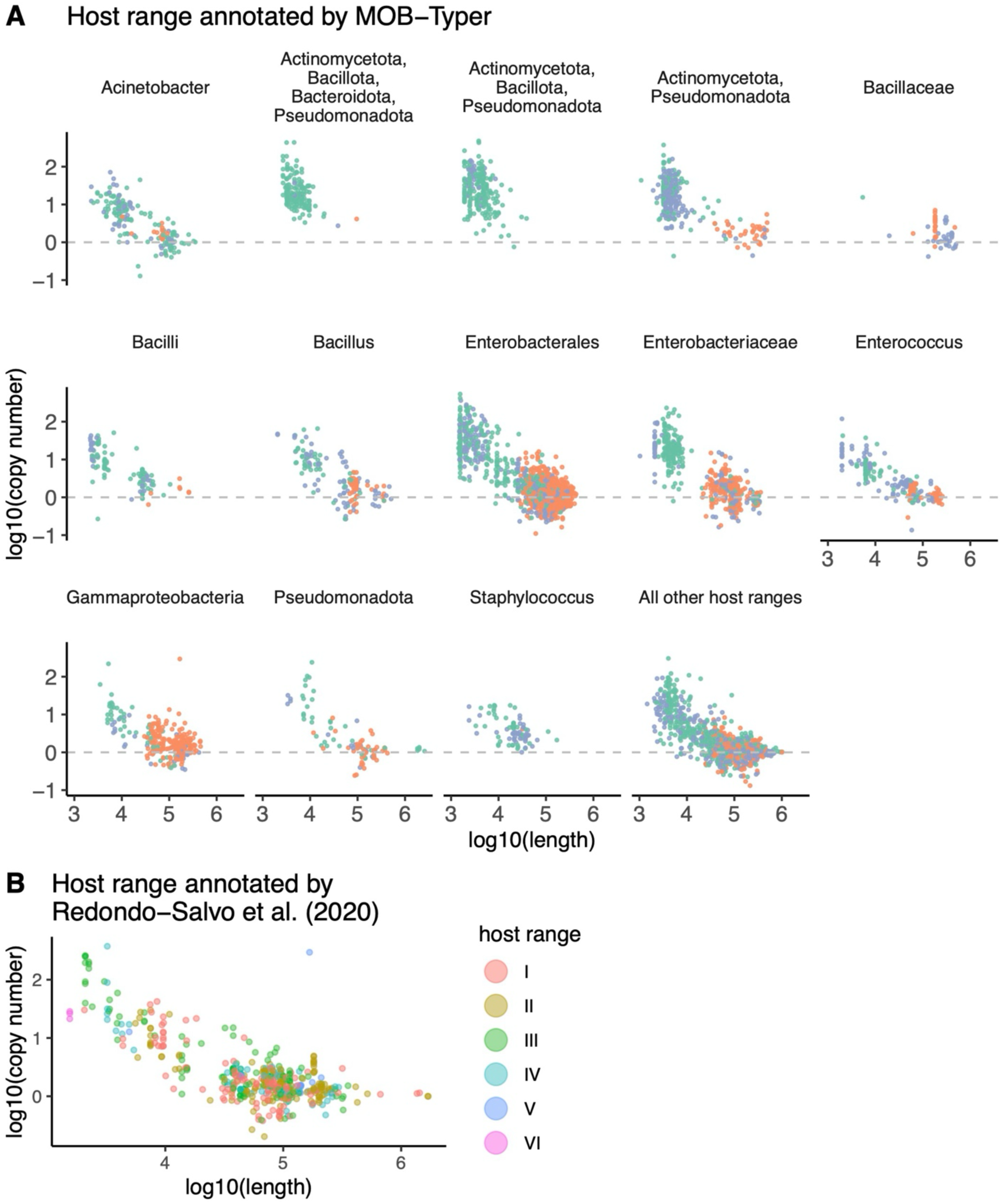
Plasmid host range does not correlate with plasmid length or copy number. Conjugative plasmids are colored orange pink, mobilizable plasmids are colored light green, and non-mobilizable plasmids are colored light blue. **A)** Host range annotated with MOB-Typer. Host ranges with more than 50 plasmids are shown in separate panels; the remaining host ranges are lumped together in the final panel. **B)** Host range annotated by Redondo-Salvo, et al. (2020). Host range are classified from I (most narrow host range) to VI (broadest host range).

**Supplementary Figure S7.**
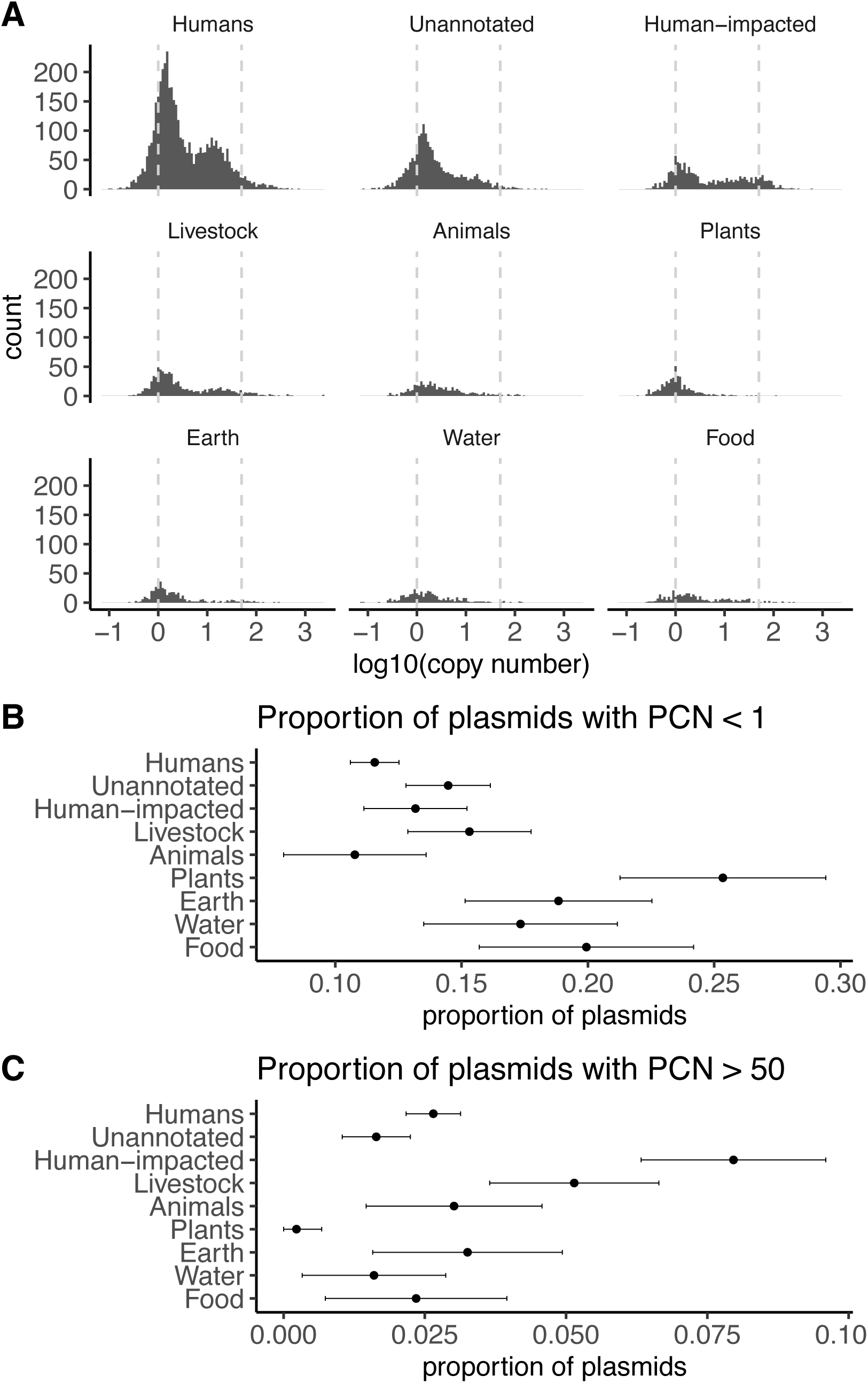
Low-copy number plasmids are common, while high copy number plasmids are rare and are enriched in human-impacted environments. **A)** Histogram of plasmid copy numbers across ecological categories. Dashed lines are drawn at PCN = 1 and PCN = 50. **B)** Proportion of very low copy number (PCN < 1) plasmids per ecological category. Each point represents the proportion of isolates containing very low copy number plasmids (PCN < 1) within each ecological category. Error bars represent 95% binomial proportion confidence intervals around the mean, using the formula 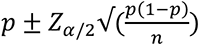, where *p* is the proportion, *n* is the sample size, and *Z_a/2_* = 1.96. **C)** Proportion of high copy number (PCN > 50) plasmids per ecological category. Each point represents the proportion of isolates containing high copy number plasmids (PCN > 50) within each ecological category. Error bars use the same formula as in panel B).

**Supplementary Figure S8.**
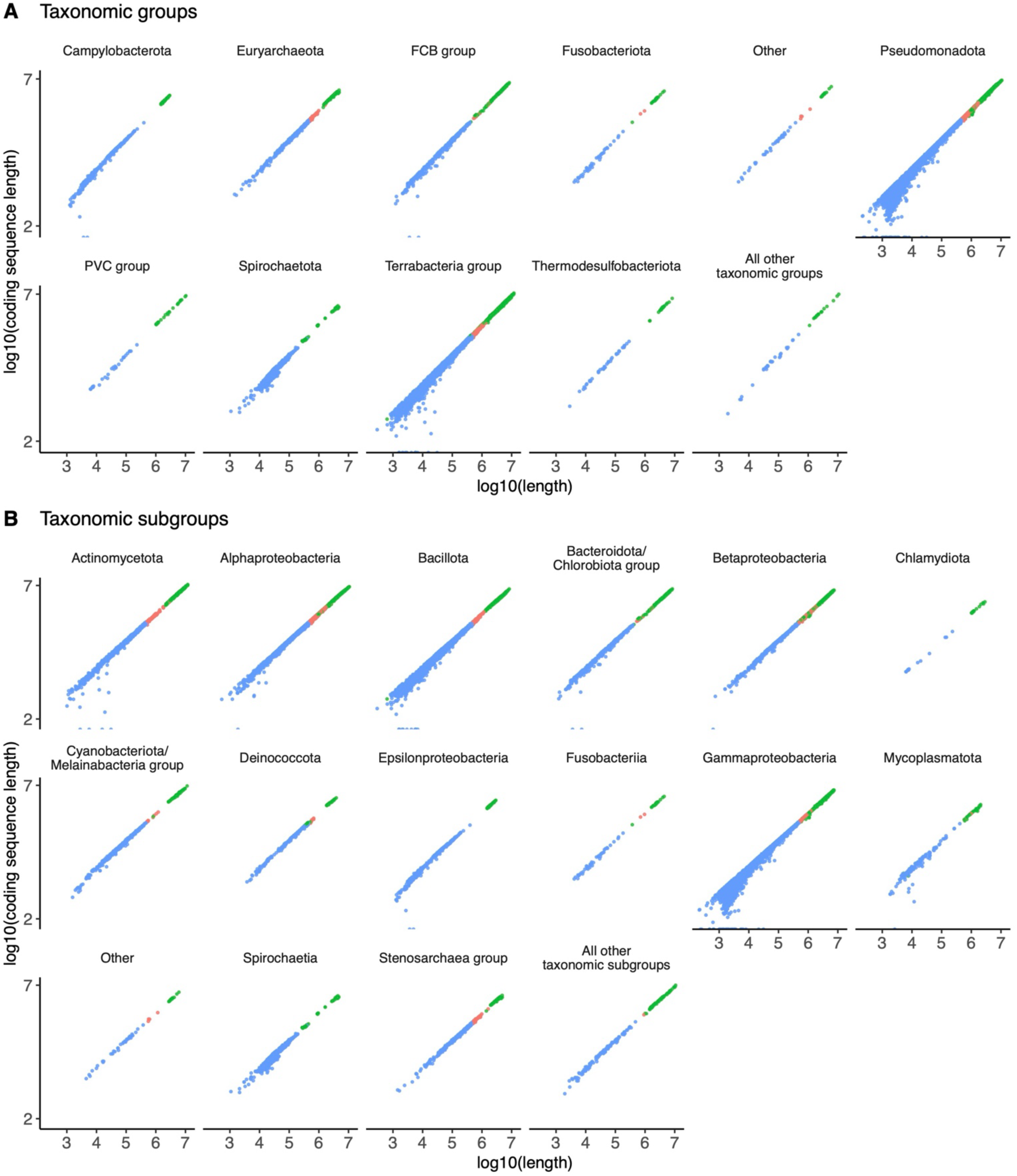
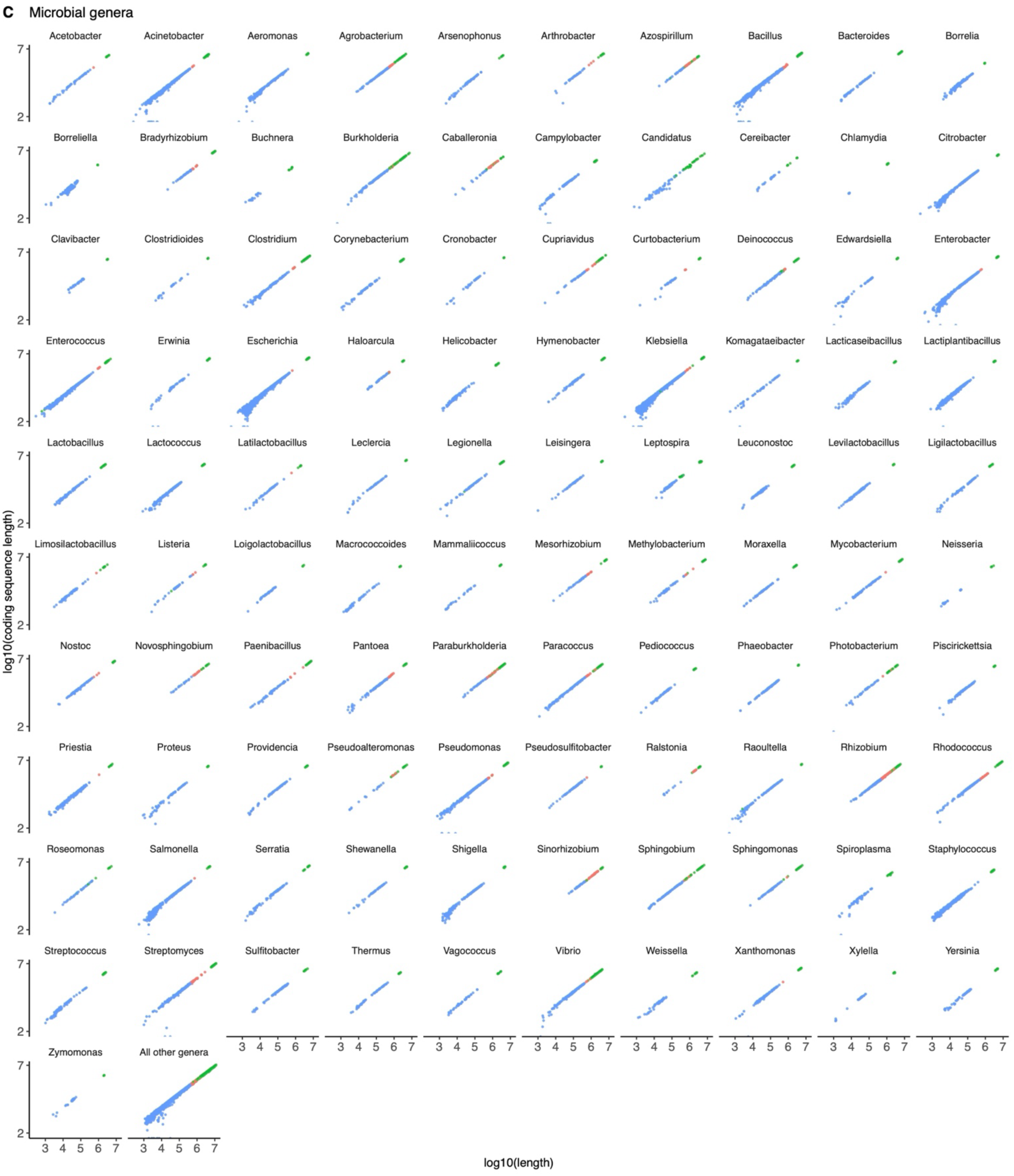
Protein-coding sequences on plasmids follow an empirical scaling law that holds across genera. Plasmids are shown in blue, megaplasmids (plasmid length > 500,000 bp) are shown in red, chromosomes are shown in green. **A)** Each panel represents a NCBI taxonomic group with at least 50 plasmids; genera with fewer than 50 plasmids are lumped together in the final panel. **B)** Each panel represents a NCBI taxonomic subgroup with at least 50 plasmids; genera with fewer than 50 plasmids are lumped together in the final panel. **C)** Each panel represents a genus with at least 50 plasmids; genera with fewer than 50 plasmids are lumped together in the final panel.

**Supplementary Figure S9.**
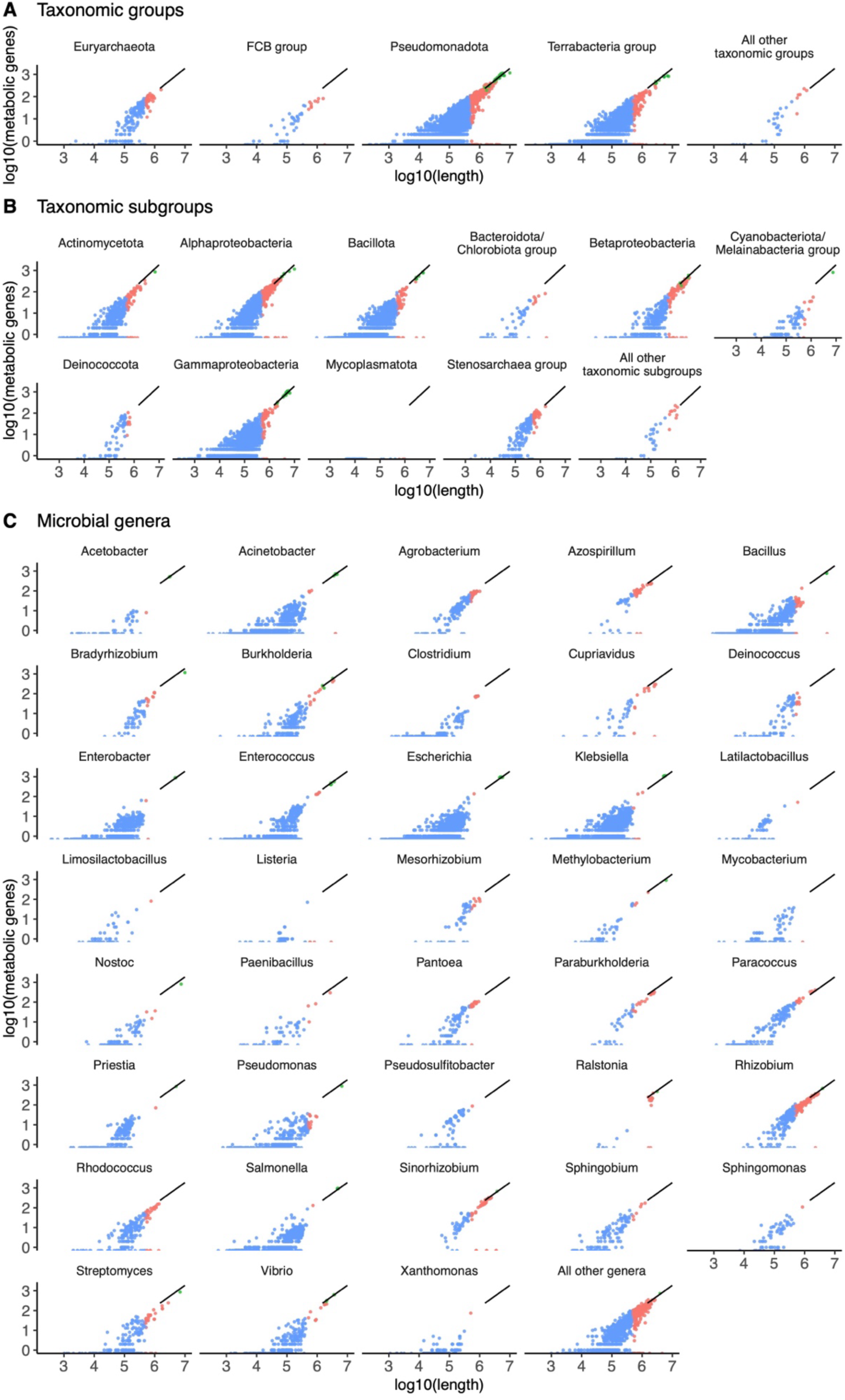
Metabolic genes on plasmids follow an empirical scaling law that holds across genera containing megaplasmids (plasmid length > 500,000 bp). Plasmids are shown in blue, megaplasmids (plasmids > 500,000 bp in length) are shown in red, chromosomes are shown in green. The linear regression between log10(metabolic genes) and log10(length) for chromosomes is shown in black. **A)** Each panel represents a NCBI taxonomic group with at least 50 plasmids; genera with fewer than 50 plasmids are lumped together in the final panel. **B)** Each panel represents a NCBI taxonomic subgroup with at least 50 plasmids; genera with fewer than 50 plasmids are lumped together in the final panel. **C)** Each panel represents a genus with at least 50 plasmids; genera with fewer than 50 plasmids are lumped together in the final panel.

**Supplementary Table 1.** Three out of fifty randomly chosen genomes containing plasmids have publications with reported plasmid copy numbers (PCN).

| Genome Accession | Plasmid copy number published | Reference URL |
| --- | --- | --- |
| GCF_000145845.2_AS<br>M14584v2 | No | https://www.frontiersin.org/journals/microbiology/articles/10.3389/fmicb.2022.826365/full |
| GCF_001690035.1_AS<br>M169003v1 | No | https://www.ncbi.nlm.nih.gov/pmc/articles/PMC4714108/ |
| GCF_002073615.2_AS<br>M207361v2 | No | https://www.nature.com/articles/s41467-019-11306-6 |
| GCF_003146705.1_AS<br>M314670v1 | No | https://www.ncbi.nlm.nih.gov/pmc/articles/PMC6535554/ |
| GCF_003193665.1_AS<br>M319366v1 | No | https://www.ncbi.nlm.nih.gov/pmc/articles/PMC5786719/ |
| GCF_003345315.1_AS<br>M334531v1 | No | https://pubmed.ncbi.nlm.nih.gov/30533938/ |
| GCF_003812945.1_AS<br>M381294v1 | No | https://www.nature.com/articles/s41467-019-11306-6 |
| GCF_004768665.1_AS<br>M476866v1 | No | https://pubmed.ncbi.nlm.nih.gov/31738764/ |
| GCF_004798785.1_AS<br>M479878v1 | No | https://pubmed.ncbi.nlm.nih.gov/31738764/ |
| GCF_009892325.1_AS<br>M989232v1 | No | https://www.frontiersin.org/journals/microbiology/articles/10.3389/fmicb.2020.00926/full AND https://www.frontiersin.org/journals/microbiology/articles/10.3389/fmicb.2020.01283/full |
| GCF_010365345.1_AS<br>M1036534v1 | No | https://www.sciencedirect.com/science/article/pii/S0269749120328050 |
| GCF_011331065.1_AS<br>M1133106v1 | No | https://academic.oup.com/jac/article/76/2/385/5961599 |
| GCF_013167675.1_AS<br>M1316767v1 | No | https://www.microbiologyresearch.org/content/journal/mgen/10.1099/mgen.0.000682 |
| GCF_013589875.1_AS<br>M1358987v1 | No | https://www.ncbi.nlm.nih.gov/pmc/articles/PMC8627209/ |
| GCF_013590775.1_AS<br>M1359077v1 | No | https://www.ncbi.nlm.nih.gov/pmc/articles/PMC8627209/ |
| GCF_013728275.1_AS<br>M1372827v1 | No | https://www.ncbi.nlm.nih.gov/pmc/articles/PMC8627209/ |
| GCF_013740875.1_AS<br>M1374087v1 | No | https://www.ncbi.nlm.nih.gov/pmc/articles/PMC8627209/ |
| GCF_014131795.1_AS<br>M1413179v1 | No | NA |
| GCF_014268495.2_AS<br>M1426849v2 | No | https://www.mdpi.com/2076-2607/9/8/1766 |
| GCF_015475575.1_AS<br>M1547557v1 | No | https://www.microbiologyresearch.org/content/journal/mgen/10.1099/mgen.0.000655 |
| GCF_016904115.1_AS<br>M1690411v1 | No | https://www.nature.com/articles/s41467-019-11306-6 |
| GCF_017165275.1_AS<br>M1716527v1 | No | https://www.microbiologyresearch.org/content/journal/mgen/10.1099/mgen.0.000682 |
| GCF_017639045.1_AS<br>M1763904v1 | No | https://pubmed.ncbi.nlm.nih.gov/33842964/ |
| GCF_017639065.1_AS<br>M1763906v1 | No | https://pubmed.ncbi.nlm.nih.gov/33842964/ |
| GCF_017639165.1_AS<br>M1763916v1 | No | https://pubmed.ncbi.nlm.nih.gov/33842964/ |
| GCF_018075145.1_AS<br>M1807514v1 | No | https://www.ncbi.nlm.nih.gov/pmc/articles/PMC8981930/ |
| GCF_018884145.1_AS<br>M1888414v1 | No | NA |
| GCF_018885165.1_AS<br>M1888516v1 | No | https://pubmed.ncbi.nlm.nih.gov/34748382/ |
| GCF_019048185.1_AS<br>M1904818v1 | No | https://www.nature.com/articles/s41467-019-11306-6 |
| GCF_019551835.1_AS<br>M1955183v1 | No | https://www.frontiersin.org/articles/10.3389/fbioe.2021.772397/full |
| GCF_020149825.1_AS<br>M2014982v1 | Yes | https://www.ncbi.nlm.nih.gov/pmc/articles/PMC9453081/ |
| GCF_021513055.1_AS<br>M2151305v1 | No | NA |
| GCF_022494825.1_AS<br>M2249482v1 | No | https://www.ncbi.nlm.nih.gov/pmc/articles/PMC9325909/ |
| GCF_022494855.1_AS<br>M2249485v1 | No | https://www.ncbi.nlm.nih.gov/pmc/articles/PMC9325909/ |
| GCF_022925255.1_AS<br>M2292525v1 | No | NA |
| GCF_023278615.1_AS<br>M2327861v1 | No | https://www.frontiersin.org/journals/microbiology/articles/10.3389/fmicb.2020.00680/full |
| GCF_024266205.1_AS<br>M2426620v1 | No | https://www.ncbi.nlm.nih.gov/pmc/articles/PMC6761502/ |
| GCF_024508075.1_AS<br>M2450807v1 | No | NA |
| GCF_024918255.1_AS<br>M2491825v1 | Yes | https://journals.asm.org/doi/full/10.1128/msystems.00476-22 |
| GCF_025349945.1_AS<br>M2534994v1 | Yes | https://journals.asm.org/doi/pdf/10.1128/mra.01012-22 |
| GCF_027944875.1_AS<br>M2794487v1 | No | https://www.biorxiv.org/content/10.1101/2022.12.08.519559v1.abstract |
| GCF_027945055.1_AS<br>M2794505v1 | No | https://www.biorxiv.org/content/10.1101/2022.12.08.519559v1.abstract |
| GCF_028335185.1_AS<br>M2833518v1 | No | https://www.frontiersin.org/journals/microbiology/articles/10.3389/fmicb.2023.1150070/full |
| GCF_029873515.1_AS<br>M2987351v1 | No | NA |
| GCF_030037835.1_AS<br>M3003783v1 | No | https://pubmed.ncbi.nlm.nih.gov/37659628/ |
| GCF_030038915.1_AS<br>M3003891v1 | No | https://pubmed.ncbi.nlm.nih.gov/37659628/ |
| GCF_900175995.1_AS<br>M90017599v1 | No | https://www.sciencedirect.com/science/article/abs/pii/S0168165617301700 |
| GCF_900620255.1_BP<br>H2986 | No | https://www.nature.com/articles/s41467-023-37200-w |
| GCF_903993065.2_AI3<br>007v1_cp | No | https://www.microbiologyresearch.org/content/journal/mgen/10.1099/mgen.0.001132 |
| GCF_905071835.1_MS<br>B1_4I | No | NA |

