## Supplementary Data 1 for "Scaling laws of plasmids across the microbial tree of life"

### Probabilistic Iterative Read Assignment (PIRA)

**Definition:** a **replicon** is general term to refer to either a chromosome or plasmid.

**Definition:** a **multiread** is a sequencing read that maps to multiple replicons (e.g., the chromosome and one or more plasmids).

**Definition:** a **uniread** is a sequencing read that maps to a unique replicon.

**PIRA initialization step:**

-- Use pseudoalignment (themisto) to assign reads to replicons.
-- make the initial PCN estimate vector $\boldsymbol{\pi}_1$ with the unireads.

$$\boldsymbol{\pi}_1 = \frac{1}{(R_1/L_1)} \begin{pmatrix} R_1/L_1 \\ R_j/L_j \\ R_n/L_n \end{pmatrix} \quad \text{where}$$

$R_j$ is the number of unireads mapping to replicon j, and $L_j$ is the length of replicon j.

We assume that the 1st replicon is the longest chromosome in the genome, with copy number 1.

-- align the multireads to the reference genome (using minimap2).
-- multireads that map to a unique location by traditional alignment are added to the set of unireads, and $\boldsymbol{\pi}_1$ is updated to include this information.

For each of the remaining multireads:

- -- get the number of matches to each replicon in the reference genome.
- -- update the match matrix **M**, where $\mathbf{M}_{ij}$ = the number of matches of multiread ***i*** to replicon ***j***.

| | replicons | | | *n* |
|---|---|---|---|---|
| | *1* | *2* | *0* | *1* |
| | *0* | *1* | *1* | *0* |
| | *1* | *1* | *0* | *0* |
| multi-reads | ... | ... | ... | ... |
| *m* | *0* | *2* | *1* | *0* |

$$= \mathbf{M} = \begin{pmatrix} 1 & 2 & 0 & 1 \\ 0 & 1 & 1 & 0 \\ 1 & 1 & 0 & 0 \\ \cdots & \cdots & \cdots & \cdots \\ 0 & 2 & 1 & 0 \end{pmatrix}$$

**PIRA iterations:**

-- Turn $\boldsymbol{\pi}_1$ into a diagonal matrix **D** = diag($\boldsymbol{\pi}_1$)
-- Weight each column of **M** by the corresponding entry of the PCN guess $\boldsymbol{\pi}_1$. We do so by multiplying **M** by **D** to make the matrix-matrix product **MD.**
-- Normalize each row of **MD** to sum to 1 (probabilistic read assignment) to make matrix **M***.
-- Sum over rows of **M*** to generate the multiread vector $\mathbf{R_D} = [\, R_{D1}\ R_{D2} \ldots R_{Dn} \,]$.

-- Update $\boldsymbol{\pi}_1$ as follows:

$$\boldsymbol{\pi}_2 = \frac{1}{(R_1 + R_{D1})/L_1} \begin{pmatrix} (R_1+R_{D1})/L_1 \\ (R_j+R_{Dj})/L_j \\ R_n+R_{Dn})/L_n \end{pmatrix}$$

Iterate until the PCN estimate vector $\boldsymbol{\pi}$ converges.
